# CENP-E initiates chromosome congression by opposing Aurora kinases to promote end-on attachments

**DOI:** 10.1101/2023.10.19.563150

**Authors:** Kruno Vukušić, Iva M. Tolić

## Abstract

Accurate cell division relies on rapid chromosome congression. The kinetochore motor protein CENP-E/kinesin-7 is uniquely required for congression of polar chromosomes. It is currently assumed that CENP-E drives congression by gliding kinetochores along microtubules independently of their biorientation. Here, by studying chromosome movement under different levels of CENP-E activity, we favor an alternative model in which CENP-E initiates congression by promoting stabilization of end-on attachments. In this way, CENP-E accelerates congression initiation without significantly contributing to subsequent movement. Stabilization of end-on attachments on polar chromosomes without CENP-E is delayed due to Aurora kinase-mediated hyperphosphorylation of microtubule-binding proteins and expansion of the fibrous corona. CENP-E counters this by reducing Aurora B-mediated phosphorylation in a BubR1-dependent manner, thereby stabilizing initial end-on attachments, facilitating removal of the fibrous corona, and triggering biorientation-dependent chromosome movement. These findings support a unified model of chromosome movement in which congression is intrinsically coupled to biorientation.

## INTRODUCTION

To ensure equal segregation into the two daughter cells, duplicated chromosomes achieve biorientation by attaching to microtubules extending from opposite spindle halves, thereby satisfying the spindle assembly checkpoint (SAC)^1–3^. During early mitosis, chromosomes move towards the equatorial region of the spindle in a process known as chromosome congression^4^. Chromosomes positioned at the center of the nucleus at the onset of mitosis rapidly achieve biorientation and are characterized by shorter congression movements, without requiring kinetochore molecular motors^4,5^. In contrast, chromosomes positioned closer to spindle poles, referred to as polar chromosomes, require the activity of the kinetochore motor protein CENtromere-associated Protein-E (CENP-E/kinesin-7) for more prominent congression movements^4^. Recent studies identified that polar chromosomes are particularly prone to alignment failure during shortened mitosis in non-transformed cells and unperturbed mitosis in transformed cells^6–8^. Thus, the congression of polar chromosomes seems to exhibit distinct dynamics, regulatory mechanisms, and final efficiency^9^.

CENP-E function in chromosome movement has been studied extensively^1,4,10–21^. The prevailing model posits that CENP-E drives the gliding of polar kinetochores, which are laterally attached to microtubules, toward the equatorial plate through its microtubule plus-end directed motor activity^22–25^. This model suggests that chromosome congression is decoupled from chromosome biorientation, which occurs near the metaphase plate^19,23^. The preference for which motor drives kinetochore movement in the tug-of-war between plus-end directed CENP-E and minus-end directed dynein is influenced by several factors. This includes a single-site phosphorylation of CENP-E at Threonine 422 (T422) by Aurora kinases near the spindle poles^26^ and the increased detyrosination of tubulin on kinetochore fibers^27^. Both modifications enhance the motor activity of CENP-E^28,29^. CENP-E is recruited to an unattached outer kinetochore by BuBR1^30^, and to a fibrous corona, which sheds from the kinetochore upon stable end-on attachment formation^15,29,31^, by the Rod-ZW10-Zwilch (RZZ) and Dynein-Dynactin complexes^30,32^.

Despite the established requirement of CENP-E for chromosome congression, several observations cannot be explained by the current models. First, polar chromosomes begin to move toward the spindle center even in the absence of CENP-E activity, but they fail to properly align and move toward the nearest spindle pole^33^, contradicting the gliding model of CENP-E function. Second, polar chromosome congression still occurs even in the complete absence of CENP-E, although it is markedly delayed^4,12,26,34,35^. The extent of similarity between CENP-E-driven congression and congression occurring independently of CENP-E remains unknown. Moreover, it remains unclear how Aurora kinases can promote chromosome congression by simultaneously activating CENP-E^26^ and inhibiting the formation of end-on attachments, a process that is reportedly also influenced by CENP-E^4,36^. As a result, the mechanisms underlying chromosome congression and the coordination of the key molecular players involved in this process remain poorly understood.

Here, we reveal that chromosome congression involves two distinct biomechanical steps— initiation and movement. CENP-E is crucial only for initiating congression by facilitating the formation or stabilization of end-on attachments, rather than directly driving chromosome movement to the spindle equator, which occurs with the same dynamics regardless of CENP-E presence or activity. We arrived at this conclusion by employing a large-scale live-cell imaging strategy using lattice light-sheet microscopy (LLSM). This technique enabled us to simultaneously observe and track numerous chromosome congression events across many cells over time and space. By combining this imaging strategy with reversible chemical inhibition, protein depletion, and expression of phospho-mutants of key congression regulators, we delineated the hierarchical molecular networks governing chromosome congression. Our data demonstrate that CENP-E initiates the stabilization of end-on attachments, a critical step in congression which precedes chromosome movement. The initiation of chromosome congression involves the formation and stabilization of end-on attachments, evidenced by super-resolution microscopy of tubulin and recruitment of the end-on attachment marker Astrin, together with a gradual reduction of SAC and fibrous corona proteins on kinetochores. Aurora kinases, as master inhibitors of congression initiation, oppose this process by destabilizing end-on attachments and promoting the expansion of the fibrous corona. CENP-E counteracts the activity of Aurora kinases in a BubR1-dependent manner, promoting rapid stabilization of end-on attachments and subsequent chromosome congression movement. In summary, we demonstrate that chromosome biorientation is coupled with chromosome congression near the spindle poles, with CENP-E serving as a crucial link between the two processes and Aurora kinases acting as the key regulatory elements.

## RESULTS

### CENP-E is essential only for the initiation of movement during chromosome congression

To unravel the molecular and mechanical basis of chromosome congression in relation to CENP-E activity^1^, we developed an assay that integrates acute protein perturbations with long-term, high-speed live-cell imaging. To simultaneously image many individual cells (1.5 mm field of view) in complete volumes (35 μm depth) with high spatiotemporal resolution (∼350 nm and 2 min/image) for a prolonged duration (>24 hours) and with minimal phototoxicity, we established a live-cell imaging approach based on lattice light-sheet microscopy (Fig. 1a; and Video 1, see also Extended Data Fig. 1 for comparison with confocal microscopy). By usage of CENP-E inactivation or depletion, referred to jointly as CENP-E perturbation, we created spindles with two distinct populations of chromosomes, those aligned at the spindle midplane and polar chromosomes, resulting in mitotic spindles with synchronized chromosomes near the spindle poles (Extended Data Fig. 1)^4^. This approach minimizes the variabilities between chromosomes typically observed in intact mitosis^37^. We combined large-scale imaging with an experimental design that allowed us to acutely vary the activity or presence of CENP-E in the human non-transformed cell line RPE-1 (Fig. 1b). Specifically, we tested the following conditions: (1) ‘CENP-E reactivated,’ achieved by acute reactivation of CENP-E motor activity via washout following inhibition with the small-molecule inhibitor GSK923295^38^; (2) ‘CENP-E depleted,’ achieved by using small interfering RNAs (siRNAs) to deplete CENP-E (Fig. 1c); (3) ‘CENP-E inhibited,’ achieved by continuous inhibition of CENP-E with GSK923295; and (4) ‘control,’ treated with DMSO (Fig. 1b). We then tracked the 3D positions of centrosomes and kinetochores over time in randomly selected cells (see Methods). Protein depletion efficiency was assessed using immunofluorescence, and was found to be high for all targets examined in this study (Extended Fig. 2a–h).

**Fig. 1.**
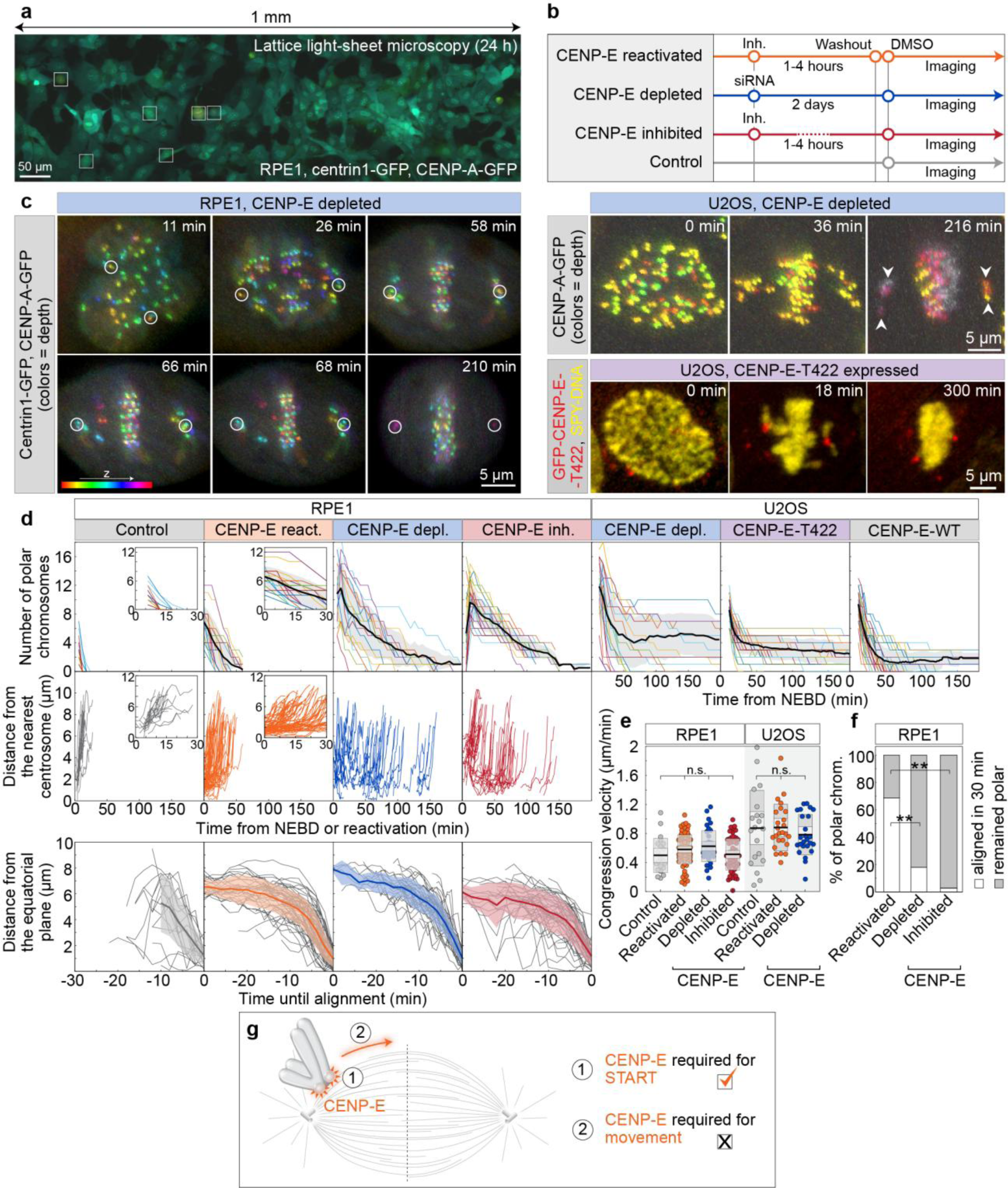
CENP-E activity is required for the rapid initiation of congression but not for the subsequent movement of polar chromosomes. **(a)** Lattice light sheet microscopy (LLSM) image of RPE-1 cells expressing centrin1–GFP and CENP-A–GFP (color coded for depth), treated with 80 nM CENP-E inhibitor (GSK-923295) for 3 h. Boxed areas highlight mitotic cells. (**b**) Schematic of the experimental approach to acutely modulate CENP-E activity. (**c**) Representative LLSM images showing: RPE-1 cell expressing centrin1–GFP and CENP-A–GFP (color coded by depth) after CENP-E depletion (left); U2OS cell with CENP-A–GFP (color coded by depth) and polar chromosomes during early anaphase (top right); and U2OS cell expressing GFP–CENP-E–T422 (red) after CENP-E depletion with SPY650–DNA staining (yellow) (bottom right). All images are maximum projections. Time 0 is mitosis onset. (**d**) Number of polar chromosomes in RPE-1 (left) and U2OS (right) cells (top), distance to the nearest spindle pole of polar kinetochore pairs in RPE-1 cells (middle) over time from NEBD or inhibitor reactivation and their distance from the equator until time of successful alignment (bottom). (**e**) Chromosome congression velocity in the final 6 min (RPE-1) or 4 min (U2OS) before alignment for indicated treatments; black line: mean, gray areas: 95% confidence interval and SD. (**f**) Percentage of polar kinetochore pairs that aligned (white) or remained polar (gray) within 30 min post-spindle elongation in RPE-1 under indicated conditions. (**g**) Diagram illustrating whether CENP-E is required for congression initiation (1) or continued movement (2), with a tick indicating the supported hypothesis. Numbers: (d and e) control (15 cells, 32 kinetochores), reactivated (18, 86), depleted (18, 69), inhibited (12, 65) (RPE-1), (d) T422 (35 cells), WT (35), depleted (21) (U2OS), (**e**): control (30 cells, 64 kinetochore pairs), reactivated (10, 38), depleted (10, 52) (U2OS), (**f**) reactivated (23 cells), depleted (38), inhibited (14) (RPE-1). All from ≥3 independent experiments. Symbols: n.s., P > 0.05; **, P ≤ 0.01; inh., inhibited; siRNA, small interfering RNA; NEBD, nuclear envelope breakdown.

**Fig. 2.**
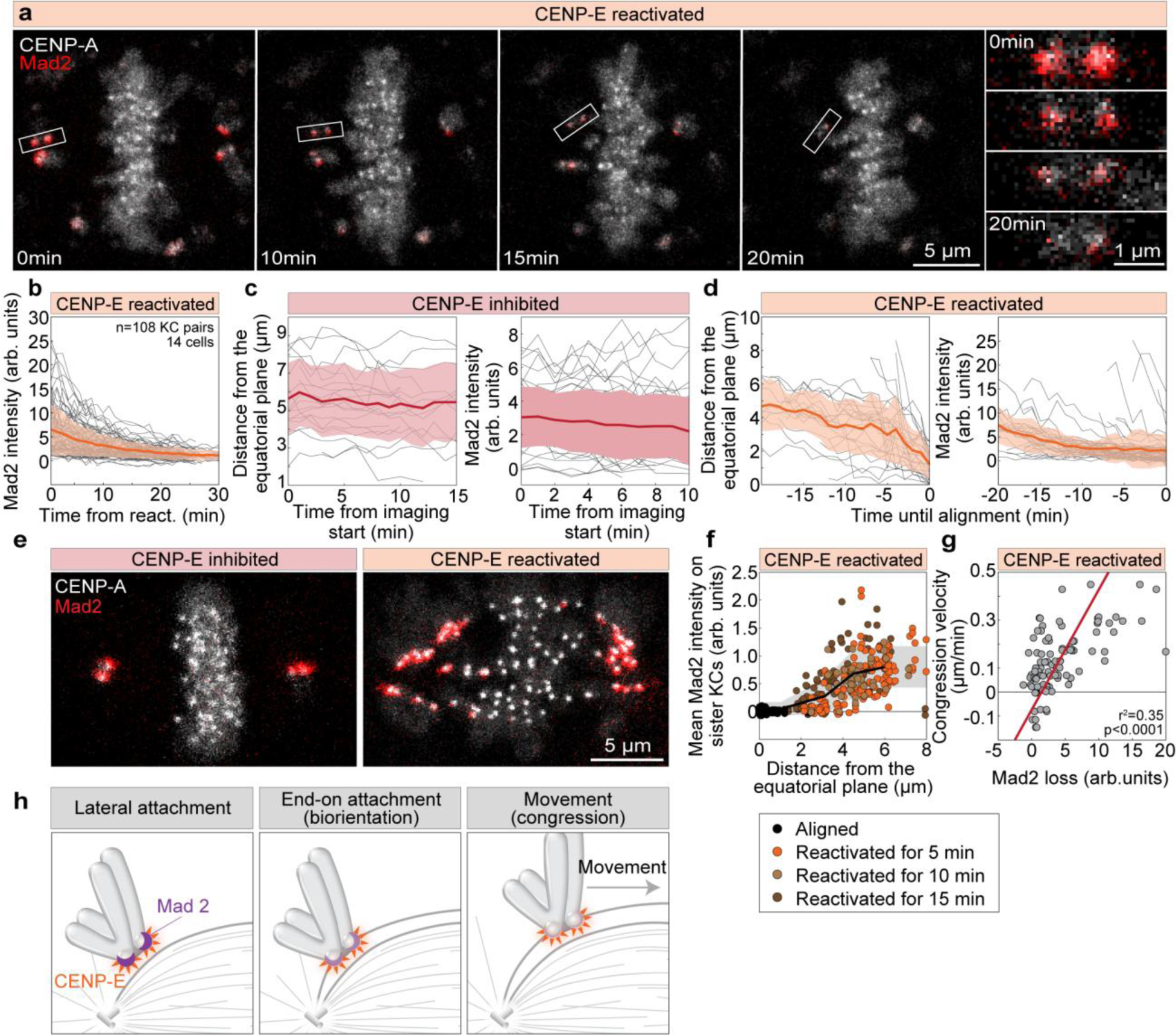
Reactivation of CENP-E triggers Mad2 loss from polar kinetochores prior to chromosome movement. **(a)** Representative time-lapse images of a cell expressing Mad2-mRuby (red) and CENPA-mCerulean (grey) after CENP-E inhibitor washout; right panels show enlarged kinetochore pairs during congression. **(b)** Mad2 intensity on initially polar sister kinetochores over time post-washout of CENP-E inhibitor, with mean (black line) and standard deviation (orange area). **(c)** Distance from equator (left) and Mad2 intensity (right) of polar kinetochores over time in presence of CENP-E inhibitor. **(d)** Distance to equator (left) and Mad2 intensity (right) of polar kinetochores over time until their alignment. **(e)** Representative images of cells fixed with or without CENP-E inhibitor showing Mad2-mRuby (red) and CENPA-mCerulean (grey). **(f)** Distance to metaphase plate versus mean Mad2 signal on kinetochores across indicated conditions; thick line: mean of binned data; grey area: standard deviation. **(g)** Congression velocity over 30 minutes after CENP-E inhibitor washout plotted against Mad2 signal loss, with regression line. Negative values reflect movement from the metaphase plate. **(h)** Diagram illustrating partial Mad2 loss (purple crescent) from kinetochores before rapid chromosome movement; CENP-E shown in orange. Complete Mad2 removal occurs progressively during congression. All images are maximum z-projections. Numbers: 14 cells, 54 kinetochore pairs (d, g); 10 cells, 28 kinetochore pairs (b); 26 cells, 135 kinetochore pairs (f); all from ≥3 biological replicates. Symbols: arb., arbitrary; inh., inhibited; depl., depleted; reac., reactivated; KC, kinetochore.

We reasoned that if chromosome congression is driven by CENP-E, the movement of polar chromosomes towards metaphase plate would differ in the presence or absence of CENP-E activity. Conversely, if CENP-E functions in processes other than directly driving chromosome movement during congression, the dynamics of polar chromosome movement should remain largely unchanged regardless of CENP-E activity. Following reactivation of CENP-E, all polar chromosomes successfully congressed to the metaphase plate during 60 min (Fig. 1d; Extended Data Fig. 1a; and Video 2). Polar kinetochores after CENP-E reactivation approached the equatorial region in a highly uniform manner, characterized by an initial slower and a faster second phase of movement (Fig. 1d; and Extended Data Fig. 1a)^19,39^, a gradual increase in interkinetochore distance (Extended Data Fig. 1b), and a gradual reorientation of the axis between sister kinetochores toward the main spindle axis (Extended Data Fig. 1c). Chromosome congression dynamics following CENP-E reactivation closely resembled those of polar kinetochores during early prometaphase in control cells (Fig. 1d; and Extended Data Fig. 1a-c; and Video 2). However, the slow phase of congression was brief or absent in control cells, as persistently polar chromosomes are absent in RPE-1 cells (Fig. 1d)^37^. We thus conclude that the CENP-E-driven congression of polar chromosomes after CENP-E reactivation recapitulates the key features of their congression during prometaphase.

How do the dynamics of chromosome congression compare between cells with the active versus perturbed CENP-E? Under CENP-E depletion or inhibition, the number of polar chromosomes gradually decreased over time as chromosomes congressed toward the metaphase plate (Fig 1d; and Video 3). Despite the absence of CENP-E or its activity, the majority of polar chromosomes successfully congressed within 3 hours, consistent with previous observations using CENP-E depletion or distinct small-molecule inhibitors^4,26,34,40–42^. Intriguingly, although polar chromosomes initiated movement asynchronously during prolonged mitosis (Fig. 1d), their movement toward the plate was strikingly similar under both active and perturbed CENP-E conditions (Fig. 1d; Extended Data Fig. 1d; Video 3). This movement was accompanied by an increase in interkinetochore distance and reorientation of the kinetochore pair toward the spindle axis in all conditions (Fig. 1d; Extended Data Fig. 1a-d). Importantly, the mean velocities of sister kinetochore movement during the 6 minutes prior to joining the plate, which corresponds to the major portion of the congression movement (Fig. 1d; Extended Data Fig. 1a-d), were indistinguishable across conditions irrespectively of CENP-E activity (Fig. 1e). This suggests that polar chromosomes move from the spindle poles to the equator in a manner independent of CENP-E activity in RPE-1 cells, challenging models in which CENP-E actively transports polar chromosomes to the metaphase plate.

Since we observed that congression movement remains highly similar regardless of CENP-E presence or activity, our next objective was to understand why the absence of CENP-E in RPE-1 cells leads to extensive delays in chromosome congression. We observed that under conditions where CENP-E was perturbed, only about 20% of polar kinetochores started congression movement within 30 minutes after spindle elongation in prometaphase. In contrast, when CENP-E was reactivated, over 85% of polar chromosomes began congression within the same 30-minute period, and 100% of them initiated congression under control conditions during this time (Fig. 1f; Extended Data Fig. 1a-d; Videos 2 and 3). This suggests that CENP-E plays a role in the initiation of kinetochore movement. Finally, we examined whether the dynamics of congression change based on the amount of time the cell spends in mitosis. There was no correlation between the velocity of congression and the time elapsed from the onset of mitosis to the initiation of congression across conditions (Extended Data Fig. 1e), indicating that the dynamics of chromosome movement during congression are robust to the time a cell spent in mitosis.

To quantify the dynamics of chromosome congression initiation in the absence of CENP-E motor activity but without directly perturbing the motor domain, we used an osteosarcoma U2OS cell line with doxycycline-inducible expression of a GFP-tagged phospho-null T422A mutant (Fig. 1c), which blocks CENP-E phosphorylation by Aurora kinases^26,43^. We compared the timing of congression initiation in these cells to that in cells expressing inducible GFP-tagged wild-type (WT) CENP-E and in CENP-E–depleted cells (Fig. 1c, Extended data Fig. 3a). Quantification of congression events over time revealed that T422A-expressing cells initiated congression of polar chromosomes similarly to CENP-E–depleted cells, whereas WT CENP-E expression enhanced congression initiation efficiency (Fig. 1d). These findings suggest that the T422A mutant functionally mimics CENP-E loss^43^, supporting the conclusion that polar chromosome initiation can proceed independently of CENP-E activity.

**Fig. 3.**
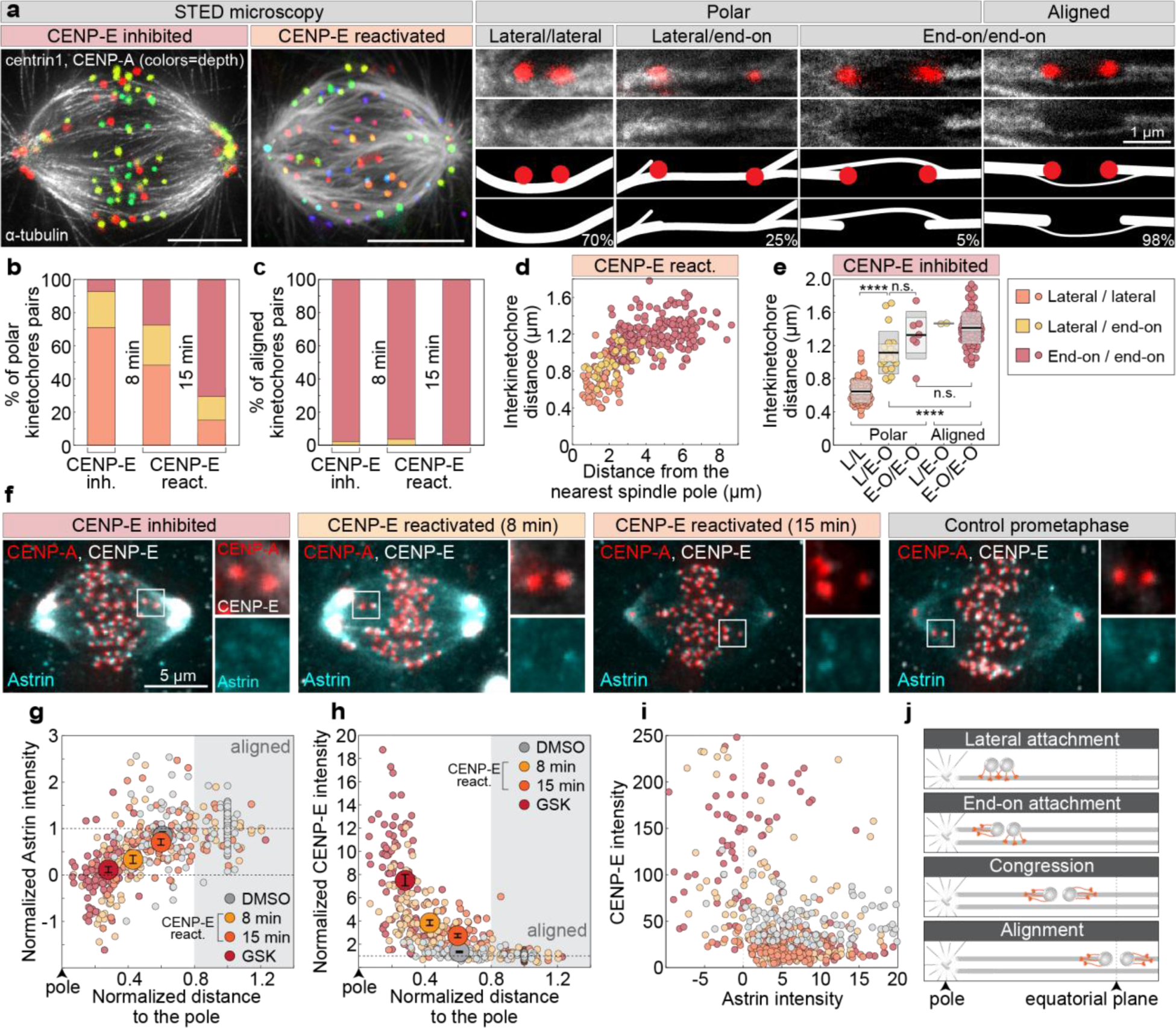
The initiation of chromosome congression near centrosomes after CENP-E reactivation coincides with chromosome biorientation. **(a)** Representative STED images of spindles in cells expressing CENP-A-GFP and centrin1-GFP (color coded for depth), either after CENP-E inhibition or 15 minutes post-reactivation, immunostained for α-tubulin (grey). Maximum projections (left) and insets showing kinetochore pairs with microtubules in different attachment categories, with corresponding diagrams (right). Percentages reflect distribution of each attachment type in CENP-E–inhibited cells. **(b, c)** Percentage of kinetochores in the indicated attachment categories for polar (b) and aligned (c) kinetochores across treatment groups (legend in e). **(d)** Interkinetochore distance versus distance from the nearest pole by attachment category (legend in e) following CENP-E reactivation. **(e)** Interkinetochore distances per attachment category in CENP-E–inhibited spindles; black line: mean; shaded areas: 95% CI. **(f)** Representative images of CENP-A–GFP and centrin1–GFP (red) cells immunostained for CENP-E (grey) and Astrin (cyan) after indicated treatments, with merged projections (left) and enlarged kinetochore views (right). **(g, h)** Astrin (g) and CENP-E (h) levels on kinetochores, normalized to CENP-A and the aligned group mean within each cell, plotted versus pole distance normalized to aligned group mean. Gray area indicates aligned region; large dots show treatment means ± SEM. **(i)** Astrin and CENP-E levels normalized to CENP-A plotted against each other across treatments. **(j)** Schematic summarizing that lateral-to-end-on attachment transition precedes major congression and occurs near spindle poles following CENP-E reactivation or in its absence. All images are maximum projections. Numbers: 69 cells, 380 kinetochore pairs (b–d); 20 cells, 185 kinetochore pairs (e); 80 cells, 505 kinetochores (f–i); all from ≥3 biological replicates. Statistics: ANOVA with post-hoc Tukey’s HSD test. Symbols: n.s., P > 0.05; ****, P ≤ 0.0001; arb., arbitrary; inh., inhibited; react., reactivated.

To examine kinetochore dynamics during congression in transformed cells, we imaged U2OS cells expressing CENP-A-GFP and measured congression velocities under untreated, CENP-E–depleted, and CENP-E–inhibited conditions (Fig. 1c, Extended data Fig. 3b, c). Consistent with our previous findings in RPE-1 cells (Fig. 1e, Extended Data Fig. 1b), both congression velocity (Fig. 1e, Extended Data Fig. 3d) and the extent of interkinetochore stretch (Extended Data Fig. 3e) during the final four minutes before alignment were similar across conditions with varying activity of CENP-E. Together, these results demonstrate that while CENP-E is not moving chromosomes during congression, it is essential for the timely initiation of congression in both transformed and non-transformed cell types (Fig. 1f).

### The speed of congression initiation is determined by the rate of end-on attachment formation

Having identified that chromosome congression movement is preceded by an initiation phase, we hypothesized that this phase requires the formation of end-on attachments at polar kinetochores. Based on this, we proposed that CENP-E actively facilitates the establishment of end-on attachments^11^ before chromosome movement begins. To test this hypothesis, we reactivated CENP-E in cells expressing the SAC protein Mad2, which accumulates on kinetochores lacking stable end-on attachments^44–46^, enabling us to track changes in kinetochore microtubule attachments following acute CENP-E reactivation. Immediately after CENP-E reactivation, polar kinetochores had high levels of Mad2, indicating the absence of end-on attachments (Fig. 2a and b; and Extended Data Fig. 4a and b)^4^. As a response to CENP-E reactivation, Mad2 levels on polar kinetochores gradually decreased over time (Fig. 2a and b; and Extended Data Fig. 4a and b, and Video 4), in contrast to the cells in which CENP-E was continuously inhibited and Mad2 levels remained constant (Fig. 2c). Interestingly, the drop of Mad2 on polar kinetochores started, on average, prior to their fast movement towards the equatorial plane although it continued to drop as kinetochores congressed (Fig. 2d; and Extended Data Fig. 4a-c). In line with that, Mad2 dropped also on kinetochores that did not initiate movement during the imaging, though at a slower rate (Extended Data Fig. 4b). These results suggest that the formation of end-on attachments at polar kinetochores depends on CENP-E activity and occurs during the initial phase of congression, prior to the rapid movement of the kinetochore toward the metaphase plate.

If end-on conversion is coupled to congression, then congressing chromosomes would predominantly arrive at the equatorial plate already bioriented. Indeed, after CENP-E reactivation, sister kinetochores closer to the spindle pole displayed higher average Mad2 signals, which decreased as most kinetochores moved toward the metaphase plate, acquiring signal intensities comparable to those of fully aligned kinetochores (Fig. 2e and f). The Mad2 signal was negatively correlated with the distance between sister kinetochores, which serves as a readout of inter-kinetochore tension (Extended Data Fig. 4d). These results suggest that after CENP-E is reactivated, polar kinetochores progressively establish more end-on attachments with microtubules, generating increased tension on the kinetochores^46^. If the extent of end-on conversion is dictating dynamics of congression, then the average congression speed over a longer period, which reflects whether or not end-on conversion occurred, is expected to be correlated with Mad2 loss. Indeed, the speed of kinetochore movement during 30 minutes after CENP-E reactivation was correlated with the rate of Mad2 loss (Fig. 2g), suggesting that the speed of congression initiation is dictated by the rate of formation of end-on attachments, which is reflected in the rate of Mad2 loss (Fig. 2h).

### Congressing chromosomes are characterized by the gradual stabilization of end-on attachments and ongoing fibrous corona stripping

To directly test if congression is coupled with biorientation, we performed super-resolution imaging of spindle microtubules by using STimulated Emission Depletion (STED) microscopy in CENP-E inhibited cells, and after reactivation of CENP-E at two time-points in RPE-1 and U2OS cells. This approach enabled us to quantify different types of microtubule attachments to kinetochores during chromosome congression (Fig. 3a; and Extended Data Fig. 4e). The reactivation of CENP-E was accompanied by chromosome congression and a gradual transition from lateral to end-on attachments on polar chromosomes in RPE-1 (Fig. 3a and b), and U2OS cells (Extended Data Fig. 4e), while the attachment status on aligned kinetochores remained unchanged (Fig. 3c). The interkinetochore distance, along with the fraction of bioriented kinetochores, progressively increased with distance from the centrosome until reaching approximately 2.5 μm, at which point nearly all polar kinetochores were bioriented (Fig. 3d). In CENP-E-inhibited cells (Extended Data Fig. 4e), congressing chromosomes situated between the metaphase plate and the spindle pole, which represented less than 5% of all polar chromosomes, were also bioriented and stretched (Fig. 3e; Extended Data Fig. 4f). Collectively, our results demonstrate that chromosome movement during congression involves conversion from lateral to end-on kinetochore-microtubule attachments close to the spindle poles, with CENP-E playing an important role in accelerating this process.

To independently test whether stabilization and maturation of end-on microtubule attachments occur at congressing chromosomes, we stained RPE-1 cells for Astrin, a positive marker of stable end-on attachments^47,48^ and CENP-E, a marker of fibrous corona^30,49^. To capture the full range of kinetochore positions from the spindle poles to the equatorial plate, we analyzed cells under CENP-E inhibition, at two time points following CENP-E reactivation, and in DMSO-treated prometaphase cells undergoing congression (Fig. 3f). We quantified the intensity of Astrin and CENP-E at all polar kinetochores and at randomly selected aligned kinetochores, together with their distance from the nearest spindle pole. Following CENP-E reactivation, Astrin levels at polar kinetochores progressively increased as kinetochores were found closer to equatorial plane, reaching values comparable to those at aligned kinetochores (Fig. 3g). Increase in Astrin on congressing kinetochores mirrored both Mad2 loss (Fig. 2f) and end-on attachment maturation observed by STED microscopy (Fig. 3d) as the distance from the pole increased. Notably, by 15 minutes post-washout, Astrin levels at congressing kinetochores became indistinguishable from those in DMSO-treated prometaphase cells (Fig. 3g). In contrast, CENP-E levels declined as kinetochores moved toward the metaphase plate across conditions (Fig. 3h). CENP-E and Astrin display a mutually exclusive localization pattern at kinetochores: kinetochores enriched in CENP-E exhibit low Astrin levels, whereas those with high Astrin show reduced CENP-E presence (Fig. 3i). These results support a model in which chromosome congression involves the progressive capture and stabilization of end-on microtubule attachments (Fig. 3j), both following CENP-E reactivation and during unperturbed mitosis. Overall, congressing chromosomes are biochemically characterized by intermediate levels of checkpoint proteins (e.g., Mad2), intermediate Astrin levels indicating the maturation of end-on attachments, and reduced levels of fibrous corona proteins like CENP-E, which reflects ongoing corona stripping.

### The expansion of the fibrous corona delays the initiation of chromosome congression in the absence of CENP-E

What inhibits the initiation of chromosome congression in the absence of CENP-E? Based on previous studies^50^, we hypothesized that the fibrous corona—a dense protein meshwork that forms on kinetochores in the absence of end-on microtubule attachments^2,51^—might play a role in this inhibition. To test if the corona inhibits congression, we compared the mean levels of CENP-E, used as corona marker, and Mad2 on kinetochores following CENP-E inhibition, CENP-E depletion (note that here ZW10 was used as a corona marker, see below), and in control cells (Fig. 4a). Our analysis revealed a strong correlation between Mad2 and CENP-E levels on polar kinetochores in both control and CENP-E-inhibited cells, which did not increase with saturating concentrations of CENP-E inhibitor (Fig. 4b and c). The average levels of both proteins on polar kinetochores were significantly higher in CENP-E-inhibited cells compared to prometaphase cells (Fig. 4b and c), indicating excessive corona expansion on polar kinetochores after CENP-E inhibition. Surprisingly, in CENP-E-depleted cells, the level of Mad2 on polar kinetochores remained similar to that in CENP-E-inhibited cells despite a significant reduction in the amount of kinetochore CENP-E (Fig. 4b and c). In line with this, using ZW10, a component of the RZZ complex and a marker of the corona^30,52^, we observed excessive recruitment of ZW10 to polar kinetochores compared to control prometaphase cells, especially following CENP-E depletion (Fig. 4d-f). In contrast, the level of the outer kinetochore marker Mis12 on polar kinetochores remained unchanged on polar kinetochores following CENP-E perturbations (Fig. 4d-f). These findings suggest that without CENP-E or its activity, corona expands on polar kinetochores (Fig. 4g), contributing to the obstruction of end-on conversion^50^ and chromosome congression.

**Fig. 4.**
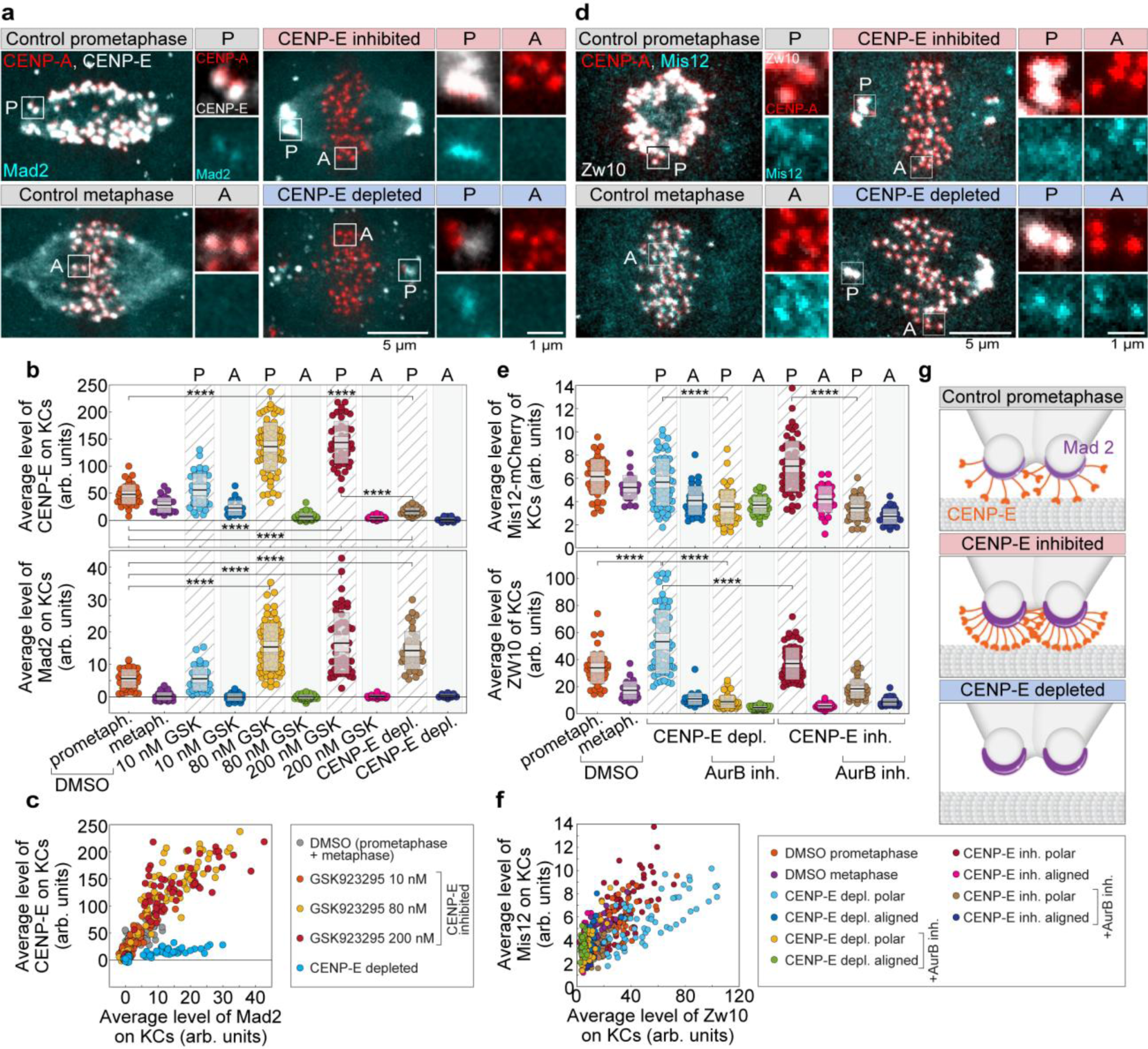
Fibrous corona expansion on kinetochores in the absence of CENP-E or its activity inhibits the initiation of chromosome congression. **(a)** Representative images of cells expressing Mad2-mRuby (cyan) and CENPA-mCerulean (red), immunostained for CENP-E (grey) after indicated treatments. Merged images (left) and enlarged views of boxed kinetochore pairs (right) are shown. A, aligned; P, polar. **(b)** Mean CENP-E (top) and Mad2-mRuby (bottom) levels on kinetochores under the indicated treatments. GSK: GSK923295. **(c)** Correlation between average Mad2 and CENP-E levels on kinetochores across treatments. **(d)** Representative cells expressing Mis12-mCherry (cyan) and CENP-A-GFP (red), immunostained for ZW10 (grey), with merged and enlarged kinetochore images. **(e)** Mean levels of Mis12-mCherry (top) and ZW10 (bottom) on kinetochores for each treatment. **(f)** Correlation between average ZW10 and Mis12 levels across treatments. All images are maximum projections. Numbers: 53 cells, 744 kinetochores (b, c); 78 cells, 810 kinetochores (e, f), all from ≥3 biological replicates. **(g)** Schematic of polar kinetochore states showing CENP-E (orange) laterally bound to microtubules (grey) and Mad2 (purple crescent) under indicated conditions. Statistics: ANOVA with post-hoc Tukey’s HSD test. Symbols: ****, P ≤ 0.0001; arb., arbitrary; inh., inhibited; depl., depleted; reac., reactivated; metaph., metaphase; prometaph., prometaphase; KC, kinetochore.

### Aurora kinases delay the initiation of congression in the absence of CENP-E

Having demonstrated that CENP-E promotes the establishment or stabilization of end-on attachments to initiate chromosome congression, we aimed to systematically explore the molecular pathways involved in inhibiting congression in the absence of CENP-E activity through acute inhibition experiments targeting various proteins (Fig. 5a). We hypothesized that, in addition to the physical barrier of the expanded fibrous corona, a molecular signaling pathway inhibits the stabilization of end-on attachments in the absence of CENP-E, thereby preventing congression. Consequently, if end-on conversion is initiated in the absence of CENP-E activity, we would expect polar chromosomes to initiate congression. To test this, we inhibited Aurora B kinase that counteracts formation of end-on attachments^36^, and maintenance of the Mad1:Mad2 as well as RZZ complex and fibrous corona on kinetochores^34,36,50,53^. Fascinatingly, following acute inhibition of Aurora B with 3 µM ZM-447439^54^ in both CENP-E-inhibited and depleted conditions, polar chromosomes immediately began moving, with most completing congression within 30 minutes after inhibitor addition (Fig. 5b-d, and Extended Data Fig. 5a-f; and Video 5). A small fraction of polar chromosomes did not reach the metaphase plate but ended their journey close to the plate (Fig. 5b, arrows; and Extended Data Fig. 5f), probably due to deficient correction of attachment errors^55^. These results imply that polar chromosomes initiate congression rapidly without CENP-E if end-on conversion is forced by the acute inhibition of Aurora B.

**Fig. 5.**
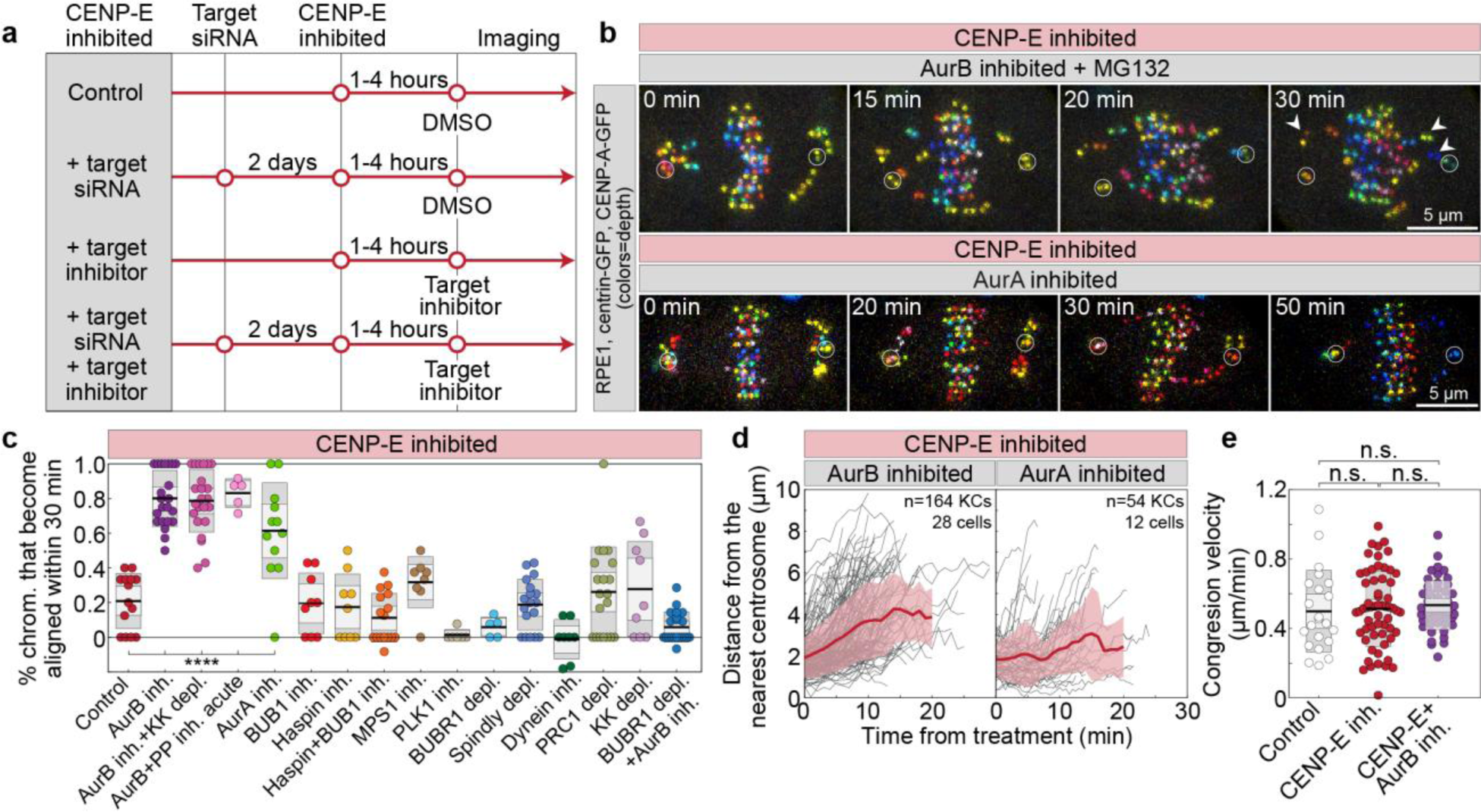
Aurora kinases inhibit initiation of chromosome congression in the absence of CENP-E. **(a)** Schematic of the experimental protocol used to test which molecular factors restrict chromosome congression when CENP-E is inhibited. **(b)** Representative images of cells expressing CENP-A-GFP and centrin1-GFP (white circles mark centrioles) following acute inhibition of target proteins, as indicated, during ongoing CENP-E inhibition. Time indicates duration post-treatment. **(c)** Percentage of polar chromosomes that aligned within 30 minutes of treatment in the presence of CENP-E inhibitor. The black line marks the mean; light and dark grey areas represent the 95% confidence interval and standard deviation, respectively. Numbers of cells per treatment: 14, 22, 23, 5, 12, 9, 9, 17, 8, 6, 5, 18, 8, 20, 9, 21, all from ≥3 biological replicates. Negative values indicate increased number of polar chromosomes after 30 minutes. **(d)** Distance of initially polar kinetochore pairs to the nearest centrosome over time after indicated treatment. Thick lines represent means; grey areas show standard deviations. **(e)** Chromosome congression velocity during the 6-minute interval preceding full alignment for control, CENP-E-inhibited, and CENP-E/Aurora B co-inhibited cells. Statistics: ANOVA with post-hoc Tukey’s HSD test. Symbols: ****, P ≤ 0.0001; Inh., inhibited; depl., depleted; siRNA, small interfering RNA; PP, protein phosphatases; KK, Kid and Kif4a; KC, kinetochore; NT, non-targeting.

Chromosome congression after acute inhibition of Aurora B in the absence of CENP-E activity was not dependent on the premature onset of anaphase, which was blocked by the proteasome inhibitor MG132^56^ (Extended Data Fig. 5g), nor on a lower concentration of ZM-447439 (Extended Data Fig. 5h) or the use of the highly specific Aurora B inhibitor Barasertib^57^ (Extended Data Fig. 5i). Furthermore, the depletion of the cohesion release factor WAPL^58^ (Extended Data Fig. 5j), the co-depletion of the chromokinesins Kid/kinesin-10 and Kif4a/kinesin-4^4^ (Fig. 5c; and Extended Data Fig. 5k), nor the acute inhibition of Protein phosphatases 1 and 2A (PP1/PP2A) by Okadaic acid^59^ (Fig. 5c; and Extended Data Fig. 5k and l) did not stop chromosomes from initiating and completing congression after acute inhibition of Aurora B. This implies, respectively, that the loss of centromeric cohesion^60^, the polar ejection forces generated by chromokinesins on chromosome arms^61,62^ and the activity of phosphatases^63^ are not essential for congression movement after acute inhibition of Aurora B in the absence of CENP-E.

Interestingly, chromosome congression in the absence of both CENP-E and Aurora B activity was marked by rapid movement of polar kinetochores toward the equatorial plane, with velocity indistinguishable from other CENP-E perturbations (Fig. 5d and e; Extended Data Fig. 5a-b and m). However, a distinctive feature of congression without Aurora B activity was the absence of the initial phase of congression, similar to control cells or, to a lesser degree, CENP-E-reactivated cells (compare Extended Data Fig. 5a and b with Fig. 1d). This suggests that only the initiation of congression is inhibited by Aurora B activity, indicating that CENP-E initiates chromosome congression by downregulating Aurora B activity.

To gain further insight into the signaling mechanisms that limit chromosome congression in the absence of CENP-E, we co-inhibited or depleted various molecular regulators involved in biorientation or Aurora B recruitment^64^ under CENP-E inhibited condition. Interestingly, acute inhibition of Aurora A kinase, which shares specificity and certain kinetochore targets with Aurora B, most notably the kinetochore protein Hec1 (Highly expressed in cancer 1)^65^ induced congression of polar chromosomes in the absence of CENP-E, although much slower than acute Aurora B inhibition (Fig. 5b-d; and Video 5). On the other hand, inhibition and co-inhibition of Bub1 (Budding uninhibited by benzimidazoles 1) or Haspin, factors essential for the recruitment of Aurora B to the outer and inner centromere, respectively^66–71^, did not induce chromosome congression in the absence of CENP-E activity, as well as inhibition of Polo-like kinase 1 (Plk1) kinase, and depletion of Bub1-related (BubR1) pseudokinase (Fig. 5c; and Extended Data Fig. 5n). The only other kinase whose inhibition showed a minor but statistically nonsignificant effect compared to the control group was Monopolar spindle 1 (Mps1) kinase (Fig. 5c; Extended Data Fig. 5n). Altogether, these results suggest that centrosome localized Aurora A and the kinetochore localized Aurora B are limiting the initiation of congression in the absence of CENP-E.

The delay in the initiation of chromosome congression in the absence of CENP-E could be due to the dominant activity of other motor proteins in the spindle. However, perturbations of microtubule motor proteins, including dynein^72–74^, as well as Kid, Kif4a, Eg5 and the crosslinker of antiparallel microtubules PRC1^75,76^ revealed that neither their indirect nor direct perturbations induced congression of polar chromosomes in the absence of CENP-E (Fig. 5c; and Extended Data Fig. 5n and o). These results suggest, respectively, that the poleward pulling forces exerted by kinetochore dynein, the activity of chromokinesins, the distance between the plus ends of microtubules nucleated in one spindle half and the opposite spindle pole, and the thickness of antiparallel microtubule bundles to which kinetochores are initially laterally attached^77^, are not major obstacles to congression in the absence of CENP-E activity. Instead, the primary barrier to congression without CENP-E is the activity of Aurora kinases.

To determine if BubR1, a known interactor of PP2A and CENP-E^36,78–80^, is required for the initiation of chromosome congression after Aurora B inhibition, we acutely inhibited Aurora B in cells lacking CENP-E and BubR1. Interestingly, in cells depleted of BubR1 and inhibited for CENP-E, the congression of polar chromosomes was completely halted following acute Aurora B inhibition, in contrast to cells with intact BubR1 (Fig. 5c). These results indicate that BubR1 is crucial for the initiation of chromosome congression, independent of its role in CENP-E recruitment^80^, and that its absence cannot be compensated for by the inhibition of Aurora kinases.

Finally, we directly tested the role of corona expansion in inhibiting congression initiation in the absence of CENP-E by quantifying ZW10 levels on polar kinetochores following CENP-E perturbations and acute Aurora B kinase inhibition. We found that acute Aurora B inhibition caused a significant decrease in ZW10 levels on polar kinetochores, reducing them to levels similar to those observed on aligned kinetochores in the absence of Aurora B inhibition, both after CENP-E inhibition and depletion (Fig. 4e, f). These findings support our conclusion that, in the absence of CENP-E or its activity, Aurora B–dependent fibrous corona expansion inhibits the initiation of chromosome congression.

### CENP-E initiates chromosome congression in a BubR1-dependent manner

We next used the acute reactivation of CENP-E, combined with the acute inhibition of other target molecules, as a tool to identify which molecular factors regulate chromosome congression both upstream and downstream of CENP-E (Fig. 6a). As a first hypothesis, we assumed that the role of CENP-E in the initiation of chromosome congression reflects the recruitment of protein phosphatases to the outer kinetochore, either directly through interaction with PP1 or through its interaction with BubR1-PP2A^26,81^. However, acute or 30-minute long inhibition (see Methods) of PP1/PP2A activities by Okadaic acid did not prevent chromosome congression after reactivation of CENP-E (Fig. 6b and c; Extended Data Fig. 6a and b; and Video 5), similarly to the finding that inhibition of PP1 in HeLa cells during prometaphase did not block congression of polar chromosomes^26^. Congression after CENP-E reactivation was also successful without the activities of Plk1 kinase which regulates microtubule dynamics and recruits PP2A to BuBR1^79,82^, Kif18a/kinesin-8, which promotes chromosome alignment at the metaphase plate^83^, or the chromokinesins Kid and Kif4a and thus the polar ejection forces they generate^61^ (Fig. 6b; and Extended Data Fig. 6c).

**Fig. 6.**
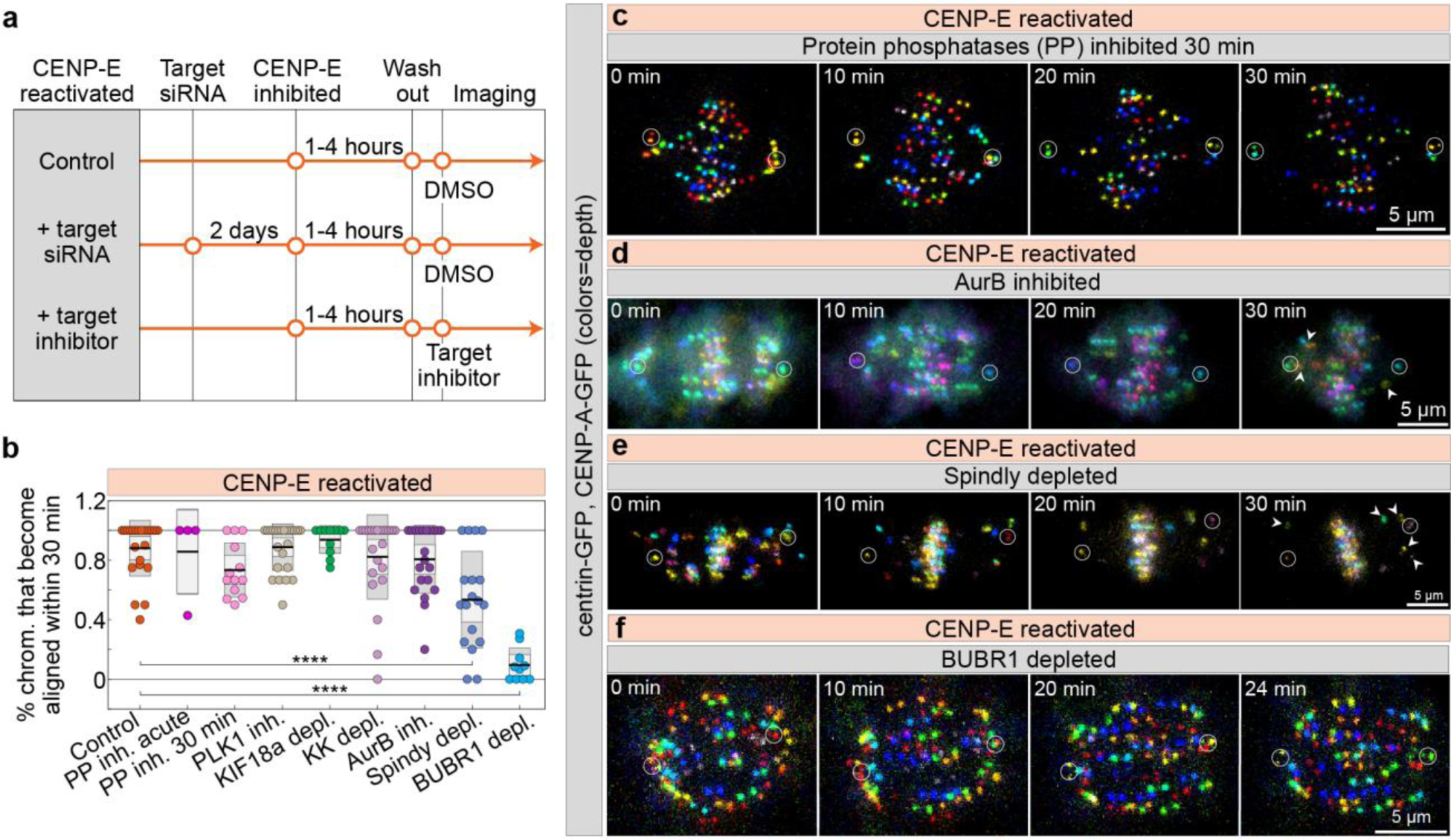
Lateral attachments and BubR1 are required for CENP-E–mediated congression initiation. **(a)** Experimental scheme testing factors regulating polar chromosome congression after CENP-E inhibitor washout. **(b, c)** Representative images of CENP-A-GFP and centrin1-GFP cells (centrioles circled) over time after washout in BubR1-depleted (b) or phosphatase-pre-inhibited (c) cells. Time indicates minutes post-washout. **(d)** Percentage of polar chromosomes aligned within 30 minutes after washout across treatments. Cell numbers: 23, 4, 13, 24, 11, 23, 21, 18, 10 per treatment, all from ≥3 biological replicates. **(e, f)** Examples of RPE1 cells expressing CENP-A-GFP and centrin1-GFP (centrioles circled) after CENP-E inhibitor GSK-923295 washout and addition of Aurora B inhibitor ZM447439 (e) or Spindly depletion (f). Images are maximum projections color-coded by depth (color bar in Fig. 1a). Arrowheads mark polar kinetochores failing to congress. Statistics: ANOVA with Tukey’s HSD post-hoc test. Symbols: ****, P ≤ 0.0001; inh., inhibited; depl., depleted; siRNA, small interfering RNA; PP, protein phosphatases; KK, Kid and Kif4a.

Phosphorylation of CENP-E by Aurora kinases has previously been reported as crucial for CENP-E’s motor activity and for the congression of polar chromosomes^26^. However, polar chromosomes were still able to congress after reactivation of CENP-E when Aurora B kinase was acutely inhibited (Fig. 6b and d). Similar to the effect of acute Aurora B inhibition when CENP-E is inactive (Fig. 5b, arrows), chromosome congression after Aurora B inhibition in the presence of CENP-E activity was occasionally accompanied by incomplete congression of a small number of polar chromosomes (Fig. 6d, arrows). This suggests that incomplete congression of polar chromosomes may result from error correction defects, such as unrepaired merotelic attachments, a known effect of Aurora B inhibition^55^. In summary, majority of polar chromosomes can initiate congression when Aurora B is fully inhibited, regardless of CENP-E activity.

It remains unclear whether CENP-E activity requires the presence of lateral attachments to mediate congression, as would be expected if CENP-E functions through the establishment of end-on attachments. To increase the number of polar chromosomes with prematurely stabilized end-on attachments, we reactivated CENP-E in cells depleted of Spindly, a dynein adaptor at the kinetochore^4,73^. The proportion of polar chromosomes that congressed after CENP-E reactivation was significantly lower in Spindly-depleted cells compared to controls (Fig. 6b, e). This suggests that CENP-E cannot initiate the congression of polar kinetochores with stable end-on attachments, or that Spindly is required to recruit CENP-E to kinetochores, as recently reported^30^.

Finally, we tested the role of BubR1 in CENP-E-mediated congression of polar chromosomes by depleting BubR1, inhibiting CENP-E, and treating cells with MG132 to prevent the premature onset of anaphase^84^. Subsequent reactivation of CENP-E activity in BubR1-depleted cells did not result in the congression of polar chromosomes, unlike in cells with intact BubR1 (Fig. 6b, f; Extended Data Fig. 6b; Video 5). This finding, along with our observation that BubR1 is required to counteract Aurora B independently of CENP-E recruitment (Fig. 5c), suggests that BubR1 is crucial for the initiation of chromosome congression.

### CENP-E initiates chromosome congression in a Hec1-dependent manner

To challenge our model that congression requires biorientation, we investigated whether CENP-E facilitates chromosome movement independently of end-on attachments in cells depleted of Hec1 and KifC1/HSET—a condition reported to permit chromosome pseudoalignment without end-on attachments^23^. We then combined this approach with CENP-E perturbations and analyzed chromosome alignment and Hec1 kinetochore levels in individual mitotic cells using immunofluorescence. Our results show that chromosome alignment improved after HSET depletion in Hec1-depleted cells, while additional CENP-E depletion led to severe misalignment (Fig. 7a, b), consistent with previous findings^23^. Notably, the effectiveness of chromosome alignment strongly correlated with residual Hec1 levels at kinetochores, which were generally higher in Hec1- and HSET-codepleted cells than in cells depleted of Hec1 alone (Fig. 7c–e; Extended Data Fig. 6d). Importantly, cells with minimal Hec1 at the kinetochores exhibited significant misalignments, regardless of HSET or CENP-E codepletion (Fig. 7a, e). We note that extensive misalignment in cells with minimal Hec1 could be exacerbated by mitotic prolongation. In live-imaged Hec1-depleted cells, the movement of polar chromosomes was unaffected by CENP-E activity (Fig. 7f), consistent with previous observations in HeLa cells^35^. These findings suggest that a low level of Hec1, about 10-15% of control levels, is sufficient to promote congression, and that CENP-E’s role in chromosome congression is promoted by the presence of Hec1.

**Fig. 7.**
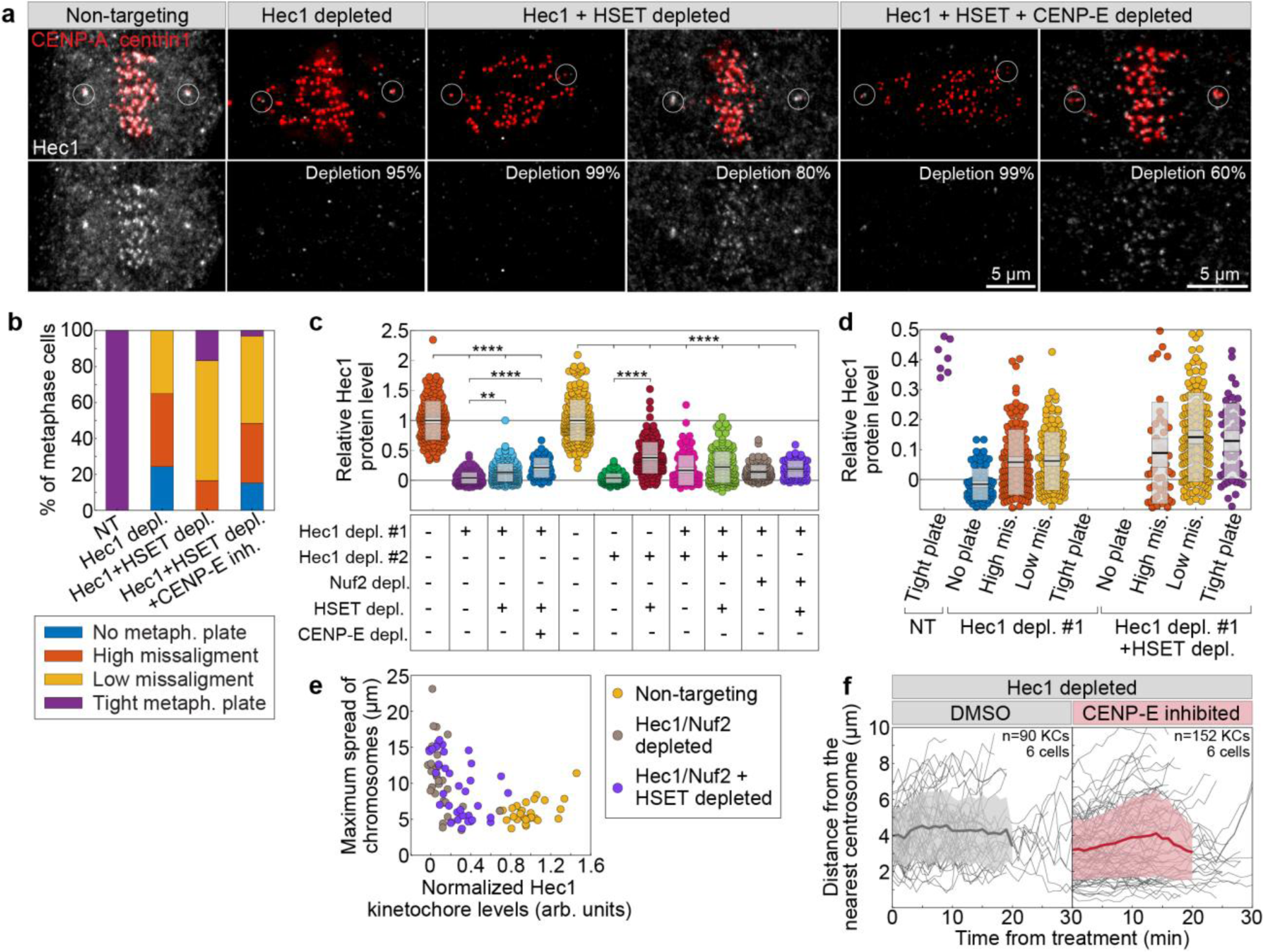
CENP-E has limited ability to initiate chromosome movement when end-on attachment formation is compromised. **(a)** Representative images of cells expressing CENP-A-GFP and centrin1-GFP (red) immunostained for Hec1 (grey) with circled centrioles, under indicated treatments. **(b)** Percentage of metaphase cells showing varying degrees of chromosome misalignment (legend) per treatments. 31, 27, 24, 33 cells per each treatment, from ≥3 replicates. **(c, d)** Average Hec1 levels on kinetochores normalized to non-targeting controls across treatments (c) and combined with alignment categories (d, legend from b). Data in (c) from 1668 kinetochores (153 cells) and in (d) from 733 kinetochores (64 cells), both from ≥3 replicates. **(e)** Hec1 levels versus maximum chromosome spread per cell across treatments. 97 cells from ≥3 replicates. **(f)** Distance over time from initially polar kinetochores to nearest centrosome under indicated treatments. All images are maximum projections. Statistics by ANOVA with Tukey’s HSD post-hoc test. Symbols: **, P ≤ 0.01; ****, P ≤ 0.0001; inh., inhibited; depl., depleted; metaph., metaphase; NT, non-targeting.

## DISCUSSION

For nearly two decades, the model that chromosomes are transported to the spindle equator without biorientation by plus-end directed kinetochore motors like CENP-E^19^ has remained widely accepted, inspiring many models of chromosome movement and regulation^4,21,23,26,27,43^. Here, we provide systematic evidence supporting an alternative model where the bulk of chromosome movement during congression depends on biorientation, consistent with earlier proposals^85,86^. Our conclusions are based on key findings: (1) chromosome movement dynamics are unaffected by CENP-E presence or activity in both transformed and non-transformed cells.; (2) end-on attachments form and the Mad2 signal decreases on polar kinetochores before their rapid movement toward the equator; (3) CENP-E levels decline early on congressing kinetochores, reflecting corona stripping that requires end-on microtubule attachments^2^; (4) all congressing chromosomes are bioriented a few microns from the centrosome, as shown by STED microscopy and Astrin labeling; (5) CENP-E has limited ability to drive chromosome movement without end-on attachments or if these attachments stabilize prematurely near the pole; (6) congression can initiate without CENP-E when Aurora kinases are inhibited, which promotes biorientation; and (7) BubR1, a known promoter of end-on conversion^87^, is essential for congression independently of its role in recruiting CENP-E.

Mechanistically, we propose that CENP-E stabilizes nascent end-on kinetochore– microtubule attachments, especially near spindle poles where high Aurora A activity likely amplifies Aurora B signaling, promoting microtubule detachment in the absence of CENP-E^88^ (Fig. 8a). This model is supported by *in vitro* evidence showing that Ndc80 and CENP-E together are sufficient for lateral-to-end-on transitions^89^. Recent studies reveal that CENP-E exists in distinct pools at the kinetochore and fibrous corona, recruited by BubR1 and the RZZ complex, respectively^30,32,80^. Given that congression initiation fully depends on BubR1, we propose that kinetochore-localized CENP-E primarily drives this process.

**Fig. 8.**
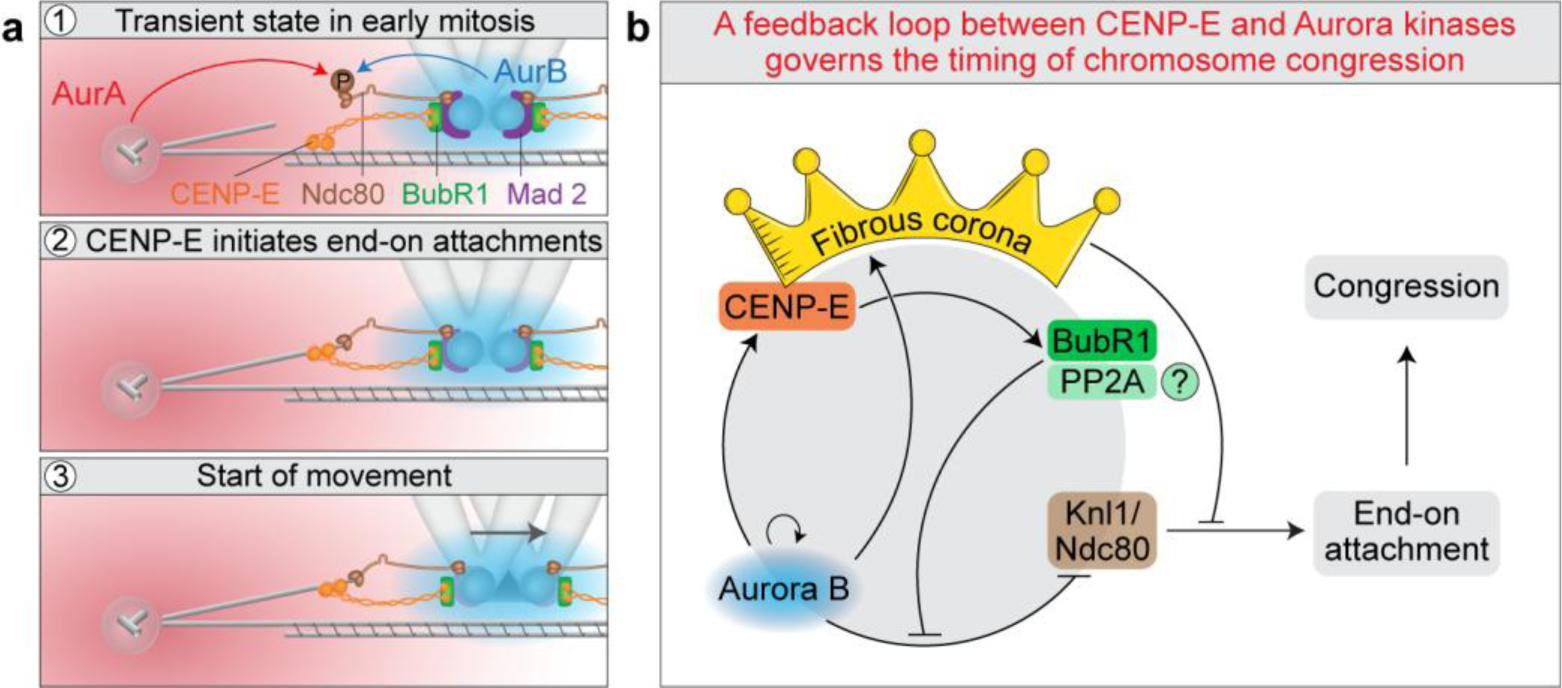
Model of chromosome congression regulation featuring a negative feedback loop between Aurora B kinase and CENP-E/BubR1. (**a**) Aurora kinases inhibit congression initiation by phosphorylating the Ndc80 tail near centrosomes (1). On polar kinetochores, CENP-E–BubR1 facilitates early end-on attachment formation near the Aurora A gradient (2), triggering a decline in Aurora B activity, loss of Mad2 from the kinetochore, and stabilization of Ndc80–microtubule binding, all preceding fast kinetochore movement (3). (**b**) This establishes a negative feedback loop involving Aurora B, CENP-E, BubR1–PP2A, the fibrous corona, and outer kinetochore proteins (Knl1 and Hec1/Ndc80). The loop self-limits by promoting end-on attachment and initiating chromosome congression, which in turn reduces the upstream signals that sustain it. Lines with arrows indicate activation, and blunt lines indicate deactivation by phosphorylation or direct physical inhibition. Initially, polar chromosomes form lateral kinetochore–microtubule attachments that fail to convert to stable end-on attachments without CENP-E due to high Aurora B activity. Aurora B phosphorylates Knl1 and Hec1 to prevent end-on conversion and maintains the expanded fibrous corona (depicted as a crown), which further inhibits attachment stabilization. Aurora B also activates CENP-E via phosphorylation. We propose (indicated by a question mark) that activated CENP-E interacts with BubR1 and possibly PP2A to counteract Aurora B-mediated phosphorylation, enabling stabilization of initial end-on attachments, progressive corona disassembly (not shown), and chromosome congression toward the spindle midplane.

We propose that the main barrier to congression initiation in the absence of CENP-E is Aurora B kinase activity (Fig. 8b). Aurora B limits the stability of end-on attachments by phosphorylating KMN network components^36,91^ and regulating the RZZ complex and fibrous corona expansion^50,53^. Aurora A also directly phosphorylates KMN near centrioles^65^. Together, Aurora kinases and the back-to-back arrangement of kinetochores^92^ prevent erroneous end-on attachments near centrosomes^4,65^, which we show hinder congression. Aurora kinases also phosphorylate and activate CENP-E near centrioles^26,43^ (Fig. 8b). We speculated that activated CENP-E interacts with BubR1–PP2A^87^, and that PP2A activity downregulates Aurora B kinase, leading to dephosphorylation of its substrates at the outer kinetochore. This stabilizes early end-on attachments on polar kinetochores, marked by the onset of Mad2 loss, thereby initiating congression (Fig. 8b). However, acute phosphatase and Plk1 inhibition, which should block phosphatase activity via BubR1^79^, did not prevent efficient congression, suggesting phosphatases may not be essential. Future studies using separation-of-function mutants of CENP-E and BubR1 could clarify this mechanism.

Aurora kinases not only phosphorylate components of the KMN network but also inhibit end-on attachment stabilization by promoting expansion of the fibrous corona on unattached chromosomes (Fig. 8b). Prior models proposed that competition between the corona—which can nucleate, sort, or capture microtubules^33,49,93^, and the Ndc80 complex regulates microtubule engagement and end-on conversion^50^. This is supported by our observations and earlier findings^48^ showing near-complete corona removal upon Aurora B inhibition, which we demonstrate triggers immediate congression. We further show that CENP-E and Astrin exhibit mutually exclusive localization at kinetochores, indicating a switch-like mechanism tied to end-on attachment formation. In cells expressing constitutively dephosphorylated Hec1, polar chromosomes only congress when CENP-E is fully depleted and not when it is inhibited^88^, indicating that hyperexpanded coronas induced by CENP-E inhibition can block congression even when KMN microtubule affinity is maximal. While disrupting Spindly or Mps1, a key corona regulators^52^, affects congression to some extent, their impact is less than that of Aurora kinase inhibition. These results suggest that KMN network phosphorylation is the main barrier to end-on attachment, with corona expansion playing a secondary role unless excessively enlarged, which can actively inhibit attachment formation.

Our model predicts that chromosome congression can initiate without CENP-E activity when the KMN network has high microtubule-binding affinity, promoting the stabilization of nascent end-on attachments and stripping of the fibrous corona (Fig. 8). We demonstrated this in polar chromosomes following Aurora kinase inhibition without CENP-E, and after its reactivation. We have also shown that a similar mechanism occurs during unperturbed mitosis, where kinetochores distant from centrosomes can stabilize end-on attachments independently of CENP-E^88^. Kinetochores unable to do so early in prometaphase are likely moved poleward by corona-associated dynein^28^ and microtubule flux^43^. In intact cells with active CENP-E, the motor likely enhances this process at all kinetochores enabling rapid biorientation and congression^33^. This is supported by findings that kinetochore fibers of aligned chromosomes are thinner when CENP-E is disrupted^85,90^. Our model also aligns with observations that constitutively dephosphorylated Hec1, which mimics high microtubule affinity, does not impair congression^91,94^, and that lateral attachments resolve even in bipolar spindles with outer kinetochore-localized Aurora B^36^.

We do not rule out that CENP-E may facilitate chromosome movement near the pole by gliding polar kinetochores along highly detyrosinated microtubules, helping them move away from the Aurora A gradient^4,27^. However, in healthy cells, chromosomes rarely approach centrosomes during mitosis^33,37^ and thus avoid high Aurora A activity—unlike in cancer cells, where polar chromosomes often move close to centrosomes early in mitosis, causing congression delays^8,95^. We propose that this spatial avoidance in healthy cells supports efficient biorientation and congression. As detailed in the accompanying manuscript^88^, centrosomal Aurora A provides a spatial cue that defines the requirement for CENP-E, which acts to counterbalance Aurora kinase activity and initiate chromosome movement.

What determines the direction of movement for polar chromosomes, causing them to migrate toward the equator rather than the spindle pole? Building on our proposed mechanism for chromosome centering during metaphase^96^, and a theoretical model of prometaphase congression^97^, we suggest that this direction is set by an asymmetry in the poleward flux of kinetochore microtubules. Specifically, kinetochore microtubules oriented toward the distant pole are expected to exhibit faster poleward flux than those facing the nearer pole, thereby producing a net force that directs chromosome movement toward the spindle center. Future research will clarify the relevance of this model to polar chromosome congression and identify the factors that control their directional movement.

The relationship between chromosome congression and biorientation has remained unclear. Earlier models proposed that congression occurs via distinct pathways—either through biorientation or through CENP-E–mediated mechanisms^1,4,19^. Our results support a model in which CENP-E links congression to biorientation near the spindle pole (Fig. 8b), reinforcing a unified view of chromosome movement. While CENP-E has been proposed to maintain lateral attachments^11^, our findings suggest it may also directly promote end-on conversion and biorientation, independently of its gliding activity. Although non-gliding roles for CENP-E have been suggested^21,98,99^, their contribution to stabilizing end-on attachments during congression remains to be tested. Our model, together with findings from the accompanying manuscript^88^, provides a molecular framework for the spatial and temporal coordination of congression and biorientation, with implications for cellular systems where these processes are highly deregulated, such as cancer cells.

## ACKNOWLEDGEMENTS

We thank Alexey Khodjakov, Jonathon Pines, and Marin Barišić for cell lines; Marin Barišić, Carlos Conde, Helder Maiato and Andrea Musacchio for reviewing and discussing the manuscript; Julie Welburn for discussions and antibodies; Magda Topić and Mia Crnogaj for help with cell culture work and preparation of inhibitors; Ivana Šarić for the drawings; and members of the Tolić group and Nenad Pavin group for constructive comments on the manuscript. This work was funded by the European Research Council (ERC Synergy Grant, GA Number 855158), the Croatian Science Foundation (HRZZ) through Swiss-Croatian Bilateral Projects (project IPCH-2022-10-9344), and projects co-financed by the Croatian Government and the European Union through the European Regional Development Fund—the Competitiveness and Cohesion Operational Programme: IPSted (Grant KK.01.1.1.04.0057) and QuantiXLie Center of Excellence (Grant KK.01.1.1.01.0004).

## AUTHOR CONTRIBUTIONS

K.V. and I.M.T conceived the project, K.V. performed all experiments, quantified, analyzed and presented the data, K.V. conceptualized and prepared original draft, K.V. and I.M.T. reviewed, edited and discussed the manuscript.

## MATERIALS & METHODS

### Reagents and Tools table

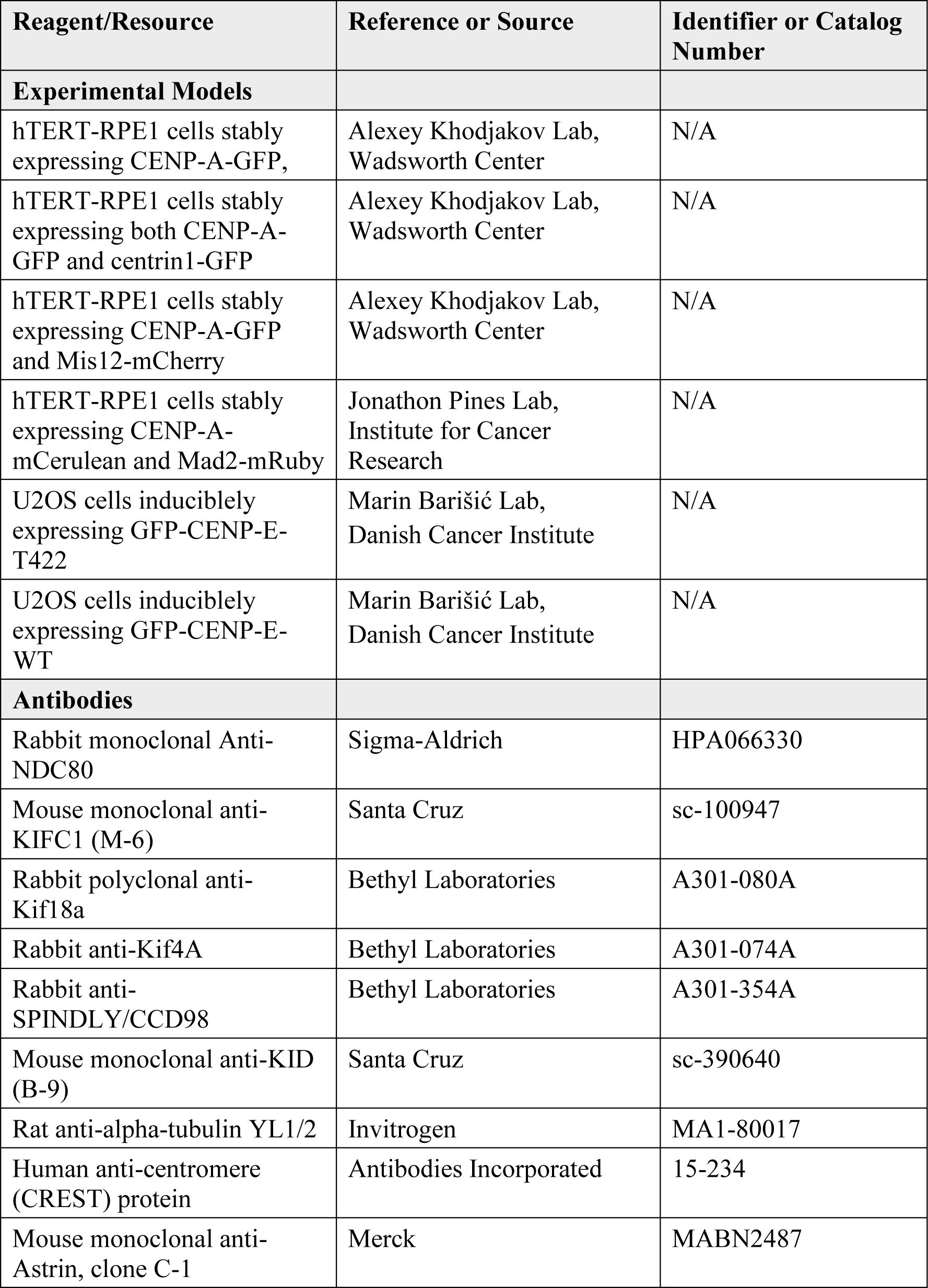
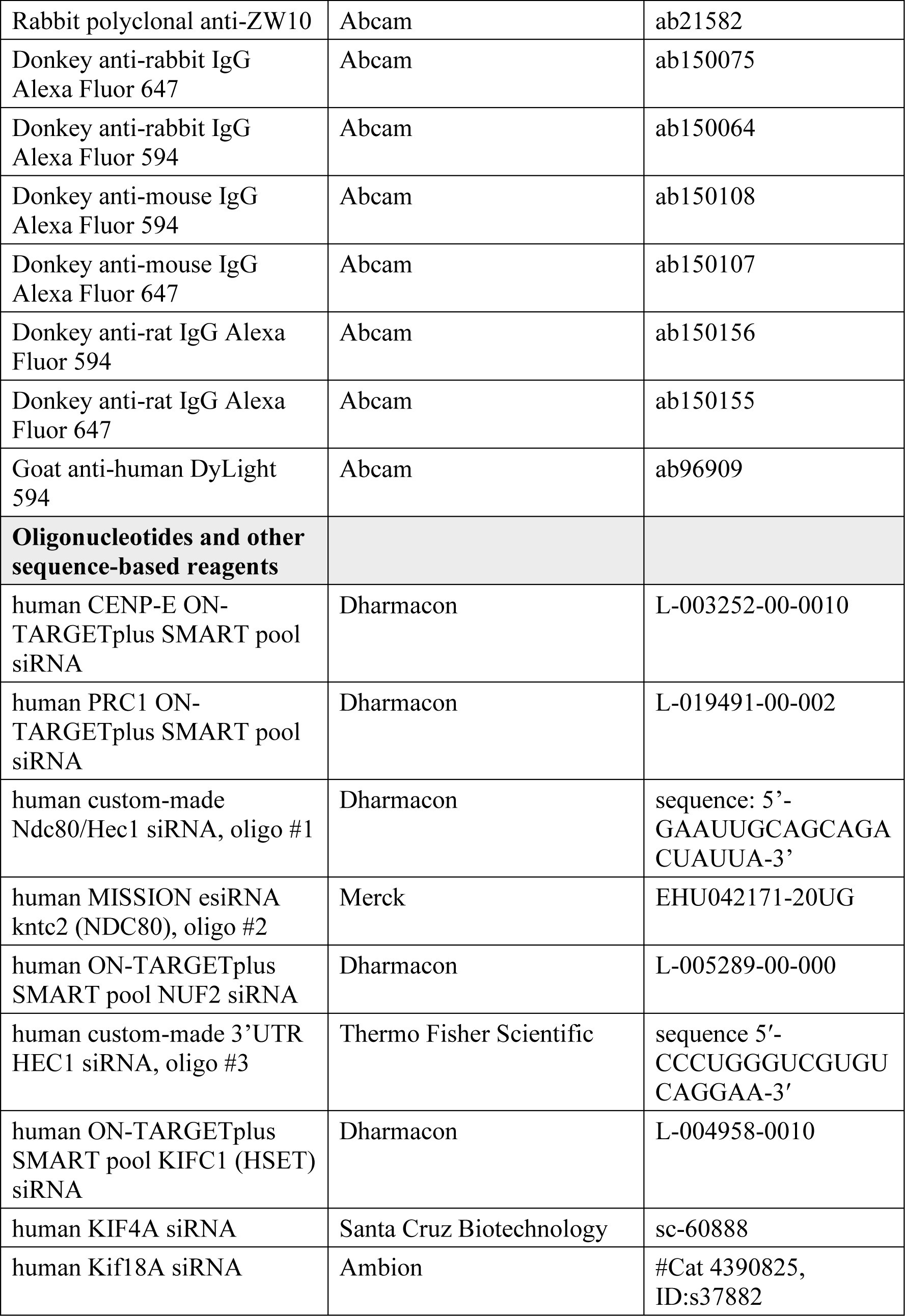
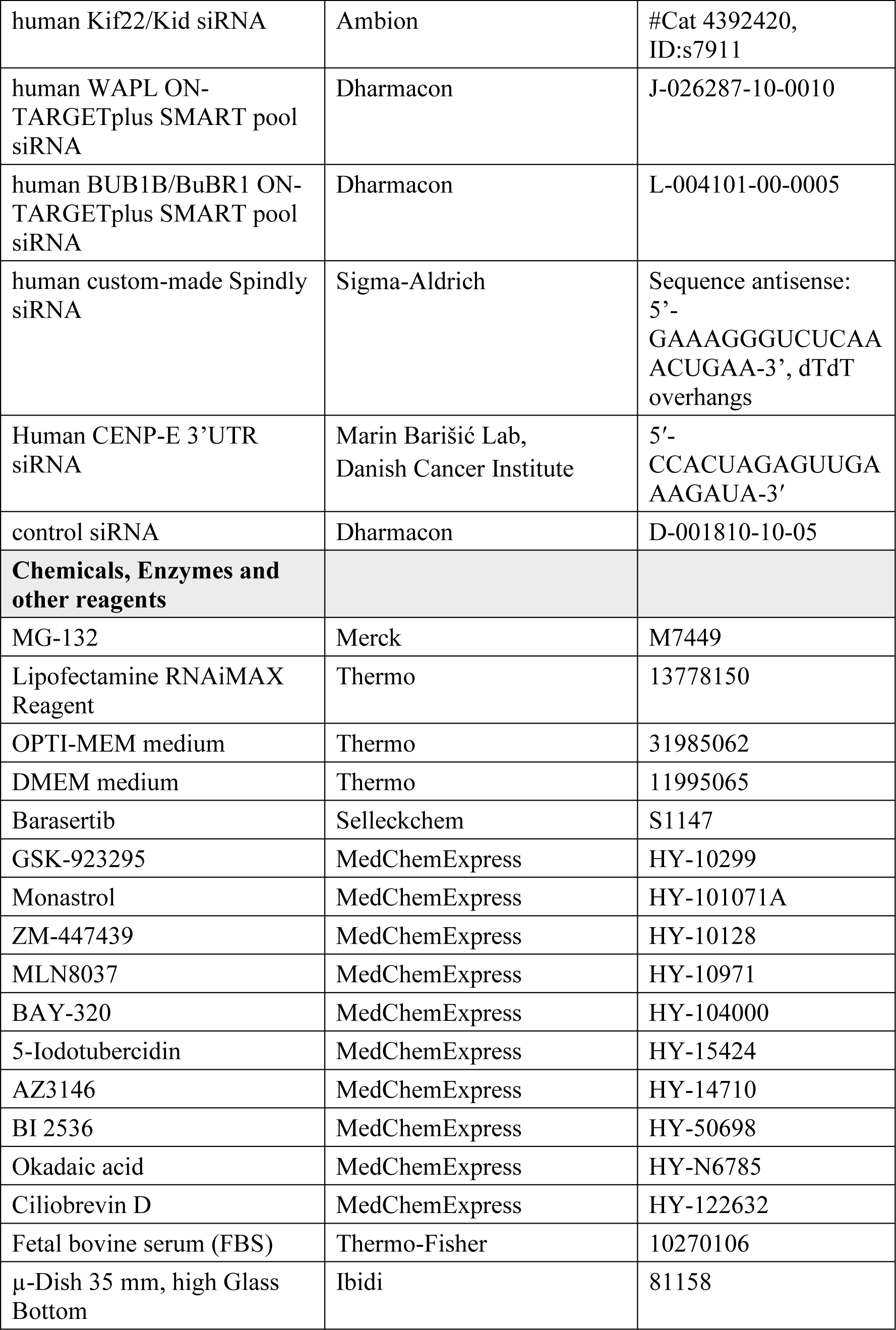
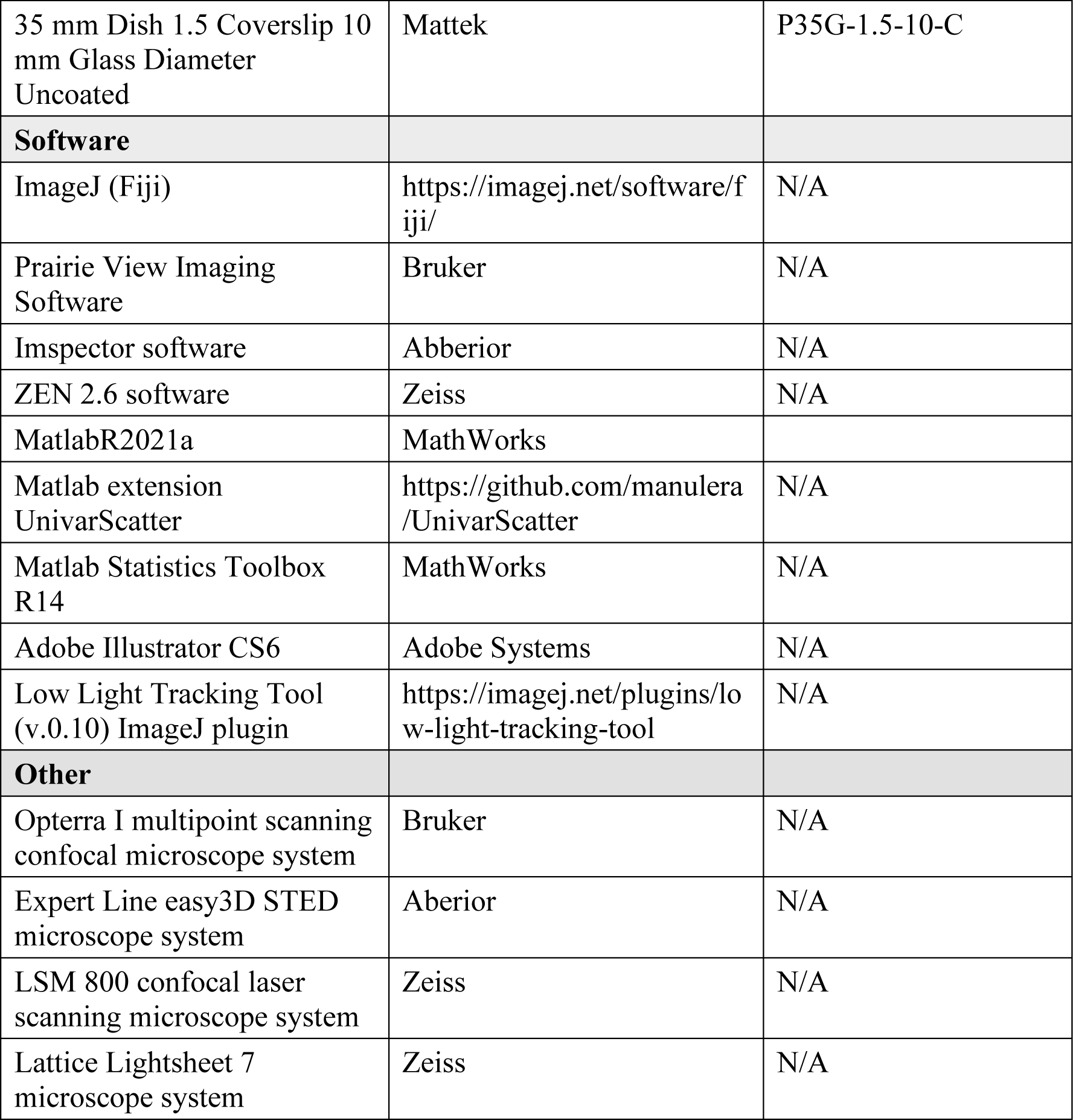

### Methods and Protocols

#### Cell lines and culture

Experiments were carried out using human hTERT-RPE1 (hTERT-immortalized retinal pigment epithelium) cells stably expressing CENP-A-GFP, human hTERT-RPE1 cells stably expressing both CENP-A-GFP and centrin1-GFP and human hTERT-RPE1 cells stably expressing CENP-A-GFP and Mis12-mCherry, all courtesy of Alexey Khodjakov (Wadsworth Center, New York State Department of Health, Albany, NY, USA), human hTERT-RPE1 cells stably expressing CENP-A-mCerulean and Mad2-mRuby, courtesy of Jonathon Pines (Institute of Cancer Research, London, UK), human HeLa (cervical carcinoma patient) cells expressing GFP-Hec1-9A, courtesy of Geert Kops (Hubrecht Institute, Utrecht, The Netherlands), human U2OS (osteosarcoma patient) cells with inducible expression of GFP-tagged wildtype (WT) CENP-E or a phosphonull CENP-E mutated at the AurA/B-specific phospho-site Threonine 422 (T422A), and human U2OS cells expressing CENP-A-GFP, were a gift from Marin Barišić (Danish Cancer Institute, Copenhagen, Denmark). All cell lines were cultured in flasks in Dulbecco’s Modified Eagle’s Medium with 1 g/L D-glucose, pyruvate and L-glutamine (DMEM, Thermo Fisher, 11995065), supplemented with 10% (vol/vol) heat-inactivated Fetal Bovine Serum (FBS, Sigma-Aldrich, St. Louis, MO, USA) and penicillin (100 IU/mL)/streptomycin (100 mg/mL) solution (Lonza, Basel, Switzerland). The cells were kept at 37 °C and 5% CO2 in a humidified incubator (Galaxy 170 S CO2, Eppendorf, Hamburg, Germany) and regularly passaged at the confluence of 70-80%. None of the cell lines were authenticated. All cell lines have also been tested for mycoplasma contamination once a month by examining samples for extracellular DNA staining with SiR-DNA (100 nM, Spirochrome, Stein am Rhein, Switzerland) and Hoechst 33342 dye (1 drop/2 ml of NucBlue Live ReadyProbes Reagent, Thermo Fisher Scientific, Waltham, MA, USA) and have been confirmed to be mycoplasma free.

#### Sample preparation and siRNAs

At 80% confluence, the DMEM medium was removed from the flask and the cells were washed with 5 ml of phosphate buffered saline (PBS). Then, 1 ml 1% trypsin/ethylenediaminetetraacetic acid (EDTA, Biochrom AG, Berlin, Germany) was added to the flask and cells were incubated at 37 °C and 5% CO2 in a humidified incubator for 5 min. After incubation, trypsin was blocked by adding 4 ml of DMEM medium. For the RNAi experiments, cells were seeded to reach 60% confluence the next day and cultured on 35 mm uncoated plates with 0.17 mm (#1.5 coverglass) glass thickness (MatTek Corporation, Ashland, MA, USA) in 1 ml of DMEM medium with the supplements described above. After one day of growth, cells were transfected with either targeting or non-targeting siRNA constructs which were diluted in OPTI-MEM medium (Thermo-Fisher) to a final concentration of 100 nM in the medium with cells. All transfections were performed 48 h before imaging using Lipofectamine RNAiMAX Reagent (Life Technologies, Waltham, MA, USA) according to the instructions provided by the manufacturer, unless otherwise indicated. Codepletions of Hec1 or Nuf2 and HSET and CENP-E in RPE-1 cells were performed for two times in two subsequent days by the same protocol. After four hours of siRNA treatment, the medium was changed to the prewarmed DMEM medium. Proteasome inhibitor MG-132 (Merck, M7449, IC50 value 100 nM) was used at a final concentration of 1 μM where indicated.

The siRNA constructs used were: human CENP-E ON-TARGETplus SMART pool siRNA (L-003252-00-0010, Dharmacon, Lafayette, CO, USA), human PRC1 ON-TARGETplus SMART pool siRNA (L-019491-00-0020, Dharmacon), human custom-made Ndc80/Hec1 siRNA (oligo #1, sequence: 5’-GAAUUGCAGCAGACUAUUA-3’, Dharmacon), human MISSION esiRNA kntc2 (NDC80) (oligo #2, EHU042171-20UG, Merck), human ON-TARGETplus SMART pool NUF2 siRNA (L-005289-00-000, Dharmacon), human ON-TARGETplus SMART pool KIFC1 (HSET) siRNA (L-004958-0010, Dharmacon), human KIF4A siRNA (sc-60888, Santa Cruz Biotechnology), human Kif18A siRNA (Ambion, #Cat 4390825, ID:s37882), human Kif22/Kid siRNA (Ambion, #Cat 4392420, ID:s7911), human WAPL ON-TARGETplus SMART pool siRNA (J-026287-10-0010, Dharmacon), human BUB1B/BuBR1 ON-TARGETplus SMART pool siRNA (L-004101-00-0005, Dharmacon, Lafayette, CO, USA), human Spindly siRNA (Sequence antisense: 5’-GAAAGGGUCUCAAACUGAA-3’, dTdT overhangs, Sigma-Aldrich, St. Louis, MO, USA), and control siRNA (D-001810-10-05, Dharmacon, Lafayette, CO, USA).

For experiments in U2OS cells expressing different CENP-E variants, endogenous CENP-E was depleted by transfecting cells with 20 nM 3′UTR-targeting siRNA (5′-CCACUAGAGUUGAAAGAUA-3′) 24 hours prior to fixation or imaging (Eibes et al., 2023). GFP-CENP-E expression was induced by adding doxycycline (1 μg/ml, Sigma-Aldrich) overnight. To label DNA, SPY650-DNA (Spirochrome) was added 6 hours before imaging together with the broad-spectrum efflux pump inhibitor verapamil (1 μM, Spirochrome). U2OS CENP-A-GFP-expressing cells were treated with CENP-E siRNA as described above for RPE-1 cells.

#### Drug treatments and drug washouts

Aurora B inhibitor Barasertib (AZD1152-HQPA, Selleckchem, Munich, Germany, IC50 value 0.32 nM) at a final concentration of 300 nM, was added acutely before imaging. CENP-E inhibitor GSK-923295 (MedChemExpress, Monmouth Junction, NJ, USA, IC50 value 3.2 nM) at a final concentration of 80 nM for RPE-1 and 200 nM for U2OS cells, to achieve the same effect, was added 1-4 hours before imaging in most experiments, or when noted acutely before imaging. Eg5 inhibitor Monastrol (HY-101071A/CS-6183, MedChemExpress, IC50 value 50 μM) at a final concentration of 100 μM, was added 3 hours before imaging or acutely before imaging when noted. Aurora B inhibitor ZM-447439 (MedChemExpress, IC50 value 130 nM) at a final concentration of 2 or 3 μM as noted, was added acutely before live imaging or 15 minutes before fixation as noted. Aurora A inhibitor MLN8237 (MedChemExpress, IC50 value 4 nM) at a final final concentration of 125 nM or 250 nM as noted, was added acutely before imaging or 30-60 minutes before fixation. Bub1 inhibitor BAY-320 (MedChemExpress, IC50 value 680 nM) at a final concentration of 10 μM, was added acutely before imaging. Haspin inhibitor 5-Iodotubercidin (5-ITu) (MedChemExpress, IC50 value 5-9 nM) at a final concentration of 2 μM, was added acutely before imaging. Mps1 inhibitor AZ3146 (MedChemExpress, USA, IC50 value 35 nM) at a final concentration of 500 nM, was added acutely before imaging. PLK1 inhibitor BI 2536 (MedChemExpress, IC50 value 0.83 nM) at a final concentration of 100 nM, was added acutely before imaging. Inhibitor of PP1 and PP2A Okadaic acid (MedChemExpress, IC50 value 0.1-0.3 nM for PP2A and 15-50 nM for PP1) at a final concentration of 1 μM, was added acutely before imaging or 30 minutes before washout of the CENP-E inhibitor as noted. Cytoplasmic dynein inhibitor Ciliobrevin D (MedChemExpress) at a final concentration of 20 μM, was added acutely before imaging. Proteasome inhibitor MG-132 (Merck, IC50 value 100 nM) at a final concentration of 1 μM, was added acutely before imaging or together with CENP-E inhibitor as noted.

The stock solutions for all drugs were prepared in DMSO except Okadaic acid which was prepared in ethanol. The stock solutions of all drugs were kept aliquoted at 10-50 μL at -20 °C for a maximum period of three months or at -80 °C for a maximum period of six months. New aliquots were thawed weekly for each new experiment. All drugs were added to DMEM media used for cell culture to obtain the final concentration of a drug as described, except Ciliobrevin D, which was added to a serum-reduced Opti-MEM medium that was placed on cells acutely before adding the drug, as Ciliobrevin D did not show any effect when used in DMEM medium with 10% FBS^74^. Drug washouts were performed by replacing drug-containing medium with 2 mL of fresh pre-warmed DMEM medium followed by four subsequent washouts with 2 mL of pre-warmed DMEM.

Acute inhibition of Aurora B by ZM-447439 (2 or 3 μM) or barasertib (300 nM) induced premature anaphase onset 15-30 min after the addition of Aurora B inhibitor, as reported previously^34^. Similarly, premature onset of anaphase was observed during the 30 min course of the imaging in cells acutely treated with the Mps1 inhibitor AZ3146 (500 nM), as expected from previous work^56^. When noted in the text, we treated CENP-E inhibited cells with the proteasome inhibitor MG-132 together with Aurora B or Mps1 inhibitors to block premature anaphase onset. MG-132 treatment provided more time for the chromosomes to complete the alignment successfully, as in the Aurora B inhibitor alone, anaphase ensued while some chromosomes were still in the alignment process.

Acute treatment of cells with dynein inhibitor 20 μM Ciliobrevin D splayed the spindle poles in all treated cells during 30 minutes after the addition of the drug, reflecting the reported role of the dynein-dynactin complex in the focus of the spindle poles in human cells^74^. Interestingly, co-inhibition of Bub1 by BAY-320 (10 μM) and Haspin by 5-ITu (2 μM) in some cells induced premature onset of anaphase during the first 20 minutes after addition of the drug, reflecting the possible role of both pathways in the maintenance of the spindle assembly checkpoint^58^. 30-minute long Okadaic acid treatment (1 μM) led to depolymerization of actin filaments and rounding of interphase cells^100^. Acute treatment of cells with Okadaic acid (1 uM) together with acute treatment with Aurora B inhibitor or after CENP-E inhibitor washout resulted in frequent expels of aligned kinetochores from the plate and sometimes complete breakage of the integrity of metaphase plates, but only after most of the polar chromosomes had already aligned to the plate, similar to previous reports of PP1 antibody injections into HeLa cells^26^. After acute treatment of CENP-E inhibited cells with the Eg5 inhibitor monastrol (50 uM), spindles quickly shortened, as expected from previous reports^101^. Similarly, acute and drastic spindle shortening was observed in most CENP-E inhibited cells after acute addition of the Aurora A inhibitor MLN8237 (125 nM), as expected from previous studies^65^. Acute inhibition of PLK1 by BI 2536 (100 nM) induced a fast block of spindle elongation and cytokinetic invasion in cells after the onset of anaphase observed only after CENP-E washout, as expected from previous reports ^102^.

Regarding cells imaged by confocal microscopy after the chronic inhibition of CENP-E by GSK-923295 (80 nM), all pseudo-metaphase spindles chosen for imaging were phenotypically similar between each other and between conditions where CENP-E was depleted or reactivated after washout of the CENP-E inhibitor: 1) in all cells few chromosomes were localized close to one of the spindle poles, called polar chromosomes, and their numbers ranged from 2-16 per cell; 2) occasionally in some cells few chromosomes were localized already in between spindle pole and equator; and 3) in all cells most chromosomes were already aligned and tightly packed at the metaphase plate. This type of chromosome arrangement was expected from previously published data that reported that only 10-30% of chromosomes upon NEBD require CENP-E-mediated alignment^4^. Likewise, under all conditions where CENP-E was perturbed and cells imaged by confocal microscopy during pseudo-metaphase, the overall displacement of spindle poles from each other was negligible (data not shown), consistent with cells being in a pseudo-metaphase state where there is no net change in spindle length. For LLSM-based assay, no inclusion criteria for imaging were used as the cells were non-synchronized and entered mitosis stochastically. Only randomly selected cells that entered mitosis and subsequently entered anaphase during imaging time were used for kinetochore and centrosome tracking analysis.

#### Immunofluorescence

All cells were fixed for 2 minutes with ice-cold methanol, except for those used in STED microscopy, which are described below. After fixation, cells were washed 3 times for 5 min with 1 ml of PBS and permeabilized with 0.5% Triton-X-100 solution in water for 30 min at room temperature. To block unspecific binding, cells were incubated in 1 ml of blocking buffer (2% bovine serum albumin, BSA) for 2 h at room temperature. The cells were then washed 3 times for 5 min with 1 ml of PBS and incubated with 500 µl of primary antibody solution overnight at 4°C. Antibody incubation was performed using a blocking solution composed of 0.1% Triton, 1% BSA in PBS.

Following primary antibodies were used: rabbit monoclonal Anti-NDC80 (HPA066330-100uL, Sigma-Aldrich, diluted 1:250), mouse monoclonal anti-KIFC1 (M-6, sc-100947, Santa Cruz, diluted 1:500), rabbit polyclonal anti-Kif18a (A301-080A, Bethyl Laboratories, diluted 1:500), mouse monoclonal anti-BubR1 (MAB3612, EMD Milipore, diluted 1:500), rabbit anti-Kif4A (A301-074A, Bethyl Laboratories, diluted 1:500), rabbit anti-SPINDLY/CCD98 (A301-354A, Bethyl Laboratories, diluted 1:500), mouse monoclonal anti-KID (B-9, sc-390640, Santa Cruz, diluted 1:500), rabbit polyclonal anti-ZW10 (ab21582, Abcam, 1:500), mouse monoclonal anti-Astrin, clone C-1 (MABN2487, Merck, 1:250), and human anti-centromere (CREST) protein (15-234, Antibodies Incorporated, CA, USA, 1:500). Where indicated, DAPI (1 µg/mL) was used for chromosome visualization.

After primary antibody, cells were washed in PBS and then incubated in 500 μL of secondary antibody solution for 1 h at room temperature. Following secondary antibodies were used: Donkey anti-rabbit IgG Alexa Fluor 594 (ab150064, Abcam, diluted 1:1000) for all rabbit antibodies, Donkey anti-mouse IgG Alexa Fluor 594 (ab150108, Abcam, diluted, 1:1000) for all mouse antibodies except anti-KID which was conjugated with Donkey anti-mouse IgG Alexa Fluor 647 (ab150107, Abcam, diluted 1:1000), goat anti-human DyLight 594 (Abcam, ab96909, diluted 1:1000), and Donkey anti-rat IgG Alexa Fluor 647 (ab150155, Abcam, diluted 1:500). Finally, cells were washed with 1 ml of PBS, 3 times for 10 min. Cells were imaged either immediately following the imaging or were kept at 4°C before imaging for a maximum period of one week.

To visualize alpha-tubulin in STED resolution in the RPE-1 CENP-A-GFP centrin1-GFP cell line, the ice-cold methanol protocol was avoided because it destroyed the unstable fraction of microtubules^103^. Cells were washed with cell extraction buffer (CEB) and fixed with 3.2% paraformaldehyde (PFA) and 0.1% glutaraldehyde (GA) in PEM buffer (0.1 M PIPES, 0.001 M MgCl2 × 6 H2O, 0.001 M EGTA, 0.5% Triton-X-100) for 10 min at room temperature. After fixation with PFA and GA, for quenching, cells were incubated in 1 ml of freshly prepared 0.1% borohydride in PBS for 7 min and then in 1 mL of 100 mM NH4Cl and 100 mM glycine in PBS for 10 min at room temperature. To block nonspecific binding of antibodies, cells were incubated in 500 μL blocking/permeabilization buffer (2% normal goat serum and 0.5% Triton-X-100 in water) for 2 h at room temperature. Cells were then incubated in 500 μL of primary antibody solution containing rat anti-alpha-tubulin YL1/2 (MA1-80017, Invitrogen, CA, SAD, diluted 1:500) overnight at 4 °C. After incubation with a primary antibody, cells were washed 3 times for 10 min with 1 ml of PBS and then incubated with 500 µl of secondary antibody containing donkey anti-rat IgG Alexa Fluor 594 (ab150156, Abcam, diluted 1:300) for 2 h at room temperature, followed by wash with 1 ml of PBS 3 times for 10 min.

#### Imaging

Opterra I multipoint scanning confocal microscope system (Bruker Nano Surfaces, Middleton, WI, USA) was mainly used for live-cell imaging of hTERT-RPE1 cells expressing CENP-A-GFP and Centrin1-GFP, in cells depleted of CENP-E by siRNAs and in cells with inhibited CENP-E by CENP-E inhibitor, in the assay we termed a confocal-based imaging assay. The system was mounted on a Nikon Ti-E inverted microscope equipped with a Nikon CFI Plan Apo VC 100x/1.4 numerical aperture oil objective (Nikon, Tokyo, Japan). The system was controlled with Prairie View Imaging Software (Bruker). During imaging, cells were kept at 37 ° C and at 5% CO2 in the Okolab cage incubator (Okolab, Pozzuoli, NA, Italy). The laser power was set to 10% for the 488nm excitation laser. For optimal resolution and signal-to-noise ratio, a 60 μm pinhole aperture was used and the xy-pixel size was set to 83 nm. For excitation of GFP, a 488-nm diode laser line was used. The excitation light was separated from the emitted fluorescence using the Opterra dichroic and Barrier 488 eGFP filter set (Chroma, USA). Images were acquired with an Evolve 512 Delta Electron Multiplying Charge Coupled Device (EMCCD) camera (Photometrics, Tucson, AZ, USA) using a 150-ms exposure time and with a camera readout mode of 20 MHz. 30-60 z-stacks encompassing all focal planes with visible green signal were acquired with 0.5 μm spacing to cover the entire spindle area in depth with unidirectional xyz scan mode and with the “fast acquisition” option enabled in software. Images of six different cells were taken simultaneously every 30s-1 min. The total duration of the time-lapse movies was 30 min to 1 hour. Movies of control U2OS cells imaged on Opterra I microscope system cells were obtained from a previous study^8^.

The STED confocal microscope system (Abberior Instruments) and the LSM800 laser scanning confocal system (Zeiss) were used for the remainder of the live-cell experiments we termed the confocal-based imaging assay and in hTERT-RPE1 cells expressing CENP-A-mCerulean and Mad2-mRuby after reactivation of CENP-E activity. STED microscopy was also used to image all fixed cells in super-resolution. STED microscopy was performed using an Expert Line easy3D STED microscope system (Abberior Instruments, Göttingen, Germany) with the 100 x/1.4NA UPLSAPO100x oil objective (Olympus, Tokio, Japan) and an avalanche photodiode detector (APD). The 488 nm line was used for excitation, with the addition of the 561 nm line for excitation and the 775 nm laser line for depletion during STED super-resolution imaging. The images were acquired using Imspector software. The xy pixel size for fixed cells was 20 nm and 10 focal planes were acquired with a 300 nm distance between the planes. For confocal live cell imaging of cells, the xy pixel size was 80 nm and 16 focal images were acquired, with 0.5 µm distance between the planes and 1 min time intervals between different frames. During imaging, live cells were kept at 37 ° C and at 5% CO2 in the Okolab stage incubation chamber system (Okolab, Pozzuoli, NA, Italy).

LSM 800 confocal laser scanning microscope system (Zeiss) was used for the rest of live-cell confocal-based imaging in hTERT-RPE1 cells expressing CENP-A-GFP and centrin1-GFP, with the following parameters: sampling in xy, 0.27 µm; z step size, 0.5 µm; total number of slices, 32; pinhole, 48.9 µm; unidirectional scan speed, 10; averaging, 2; 63x Oil DIC f/ELYRA objective (1.4 NA), 488 nm laser line (0.1-1% power for different experiments), and detection ranges of 450–558 nm for the green channel, 561 nm laser line (0.1-1% power for different experiments) and detection range of 565-650 nm for the red channel, 640 nm laser line (0.1-1% power for different experiments), and detection range of 656-700 nm for the far red channel, and 405 nm laser line (0.5% power) and detection range of 400-450 for the blue channel. Images were acquired using ZEN 2.6 (blue edition; Zeiss). Six cells in volumes of 15 µm were imaged sequentially every 1 minute at different positions determined right after the addition of a drug. During imaging, cells were incubated at 37 °C and 5% CO2 using a Pecon stage incubation chamber system (Pecon, Heating Insert P S1, #130-800 005). The system was also used for imaging of all other fixed cells that were not imaged by STED microscopy, on multiples positions and at the confocal lateral resolution.

The Lattice Lightsheet 7 microscope system (Zeiss) was used for live cell imaging of hTERT-RPE1 cells expressing CENP-A-GFP and Centrin1-GFP and U2OS cells expressing CENP-A-GFP after reactivation of CENP-E activity, in cells depleted of CENP-E by siRNAs in cells with inhibited CENP-E by CENP-E inhibitor, and in U2OS cells expressing CENP-E mutants, in the assay we called LLSM-based imaging assay. The system was equipped with an illumination objective lens 13.3×/0.4 (at a 30° angle to cover the glass) with a static phase element and a detection objective lens 44.83×/1.0 (at a 60° angle to cover the glass) with an Alvarez manipulator. Images were acquired using ZEN 2.6 (blue edition; Zeiss). The automatic immersion of water was applied from the motorized dispenser at an interval of 20 or 30 minutes. Right after sample mounting, four steps of the ‘create immersion’ auto immersion option were applied. The sample was illuminated with a 488-nm diode laser (power output 10 Mw) with laser power set to 1-2%. The detection module consisted of a Hamamatsu ORCA-Fusion sCMOS camera with exposure time set to 15-20 ms. The LBF 405/488/561/642 emission filter was used. During imaging, cells were kept at 37 °C and at 5% CO2 in a Zeiss stage incubation chamber system (Zeiss). The width of the imaging area in the x dimension was set from 1 to 1.5 mm.

#### Tracking of centrosomes and kinetochores

The spatial x and y coordinates of the kinetochore pairs were extracted in each time frame using the Low Light Tracking Tool (v.0.10), an ImageJ plugin based on the Nested Maximum Likelihood Algorithm, as previously described^101^. Tracking of polar kinetochores in the x and y planes was performed on the maximum intensity projections of all acquired z planes. Some kinetochore pairs could not be successfully tracked in all frames, mainly owing to cell and spindle movements in the z-direction over time. Spindle poles were manually tracked with points placed between the center of the two centrioles or centriole in centrinone treated cells. Kinetochore pairs that were aligned at the start of the imaging in a confocal-based imaging assay were also manually tracked in two dimensions. Quantitative analysis of all parameters was performed using custom-made MATLAB scripts (MatlabR2021a 9.10.0) scripts.

Due to the inability to reliably track polar kinetochores across all time frames due to neighboring kinetochores, tracking of polar kinetochores routinely commenced a few frames before the kinetochore pair began moving towards the equatorial plane. In the CENP-E reactivation assay, kinetochores were tracked from the onset of imaging until the successful identification of the same kinetochore pair, which occurred when the kinetochore pair was lost in the imaging plane, among other kinetochores, or when the imaging session ended, whichever came first. Aligned kinetochore pairs were tracked in the confocal-based assay using the same protocol.

#### Quantification of mean Mad2 intensity of kinetochore pairs

The fluorescence intensity signal of each kinetochore in both CENP-A and Mad2 was measured by using the “*Oval selection*” tool in ImageJ with the size and position defined by the borders of the CENP-A signal of each kinetochore in the sum intensity projection of all acquired z-planes. The background fluorescence intensity measured in the cytoplasm was subtracted from the obtained values, and the calculated integrated intensity value was divided by the number of z-stacks used to generate the sum projection of each cell. The obtained mean intensity value subtracted for background for each kinetochore was normalized to the intensity of the CENP-A signal.

#### Quantification of mean signal intensities of proteins after RNAi interreference

The fluorescence intensity signals were measured in triplicate samples for targeting and non-targeting groups for each siRNA treatment. Each siRNA targeting and non-targeting triplicates were imaged by the same protocol. All proteins were measured in mitotic cells at their respective locations during mitosis by using the “*Polygon selection tool*”, as established by many previous studies: kinetochores and chromosomes defined by the DAPI or CENP-A area on sum projection for CENP-E, Spindly, BubR1, and on single planes for Hec1 defined by the CENP-A signal; entire spindle region defined by extent of centrin-1 and CENP-A signals on sum projections for Kif4a, Kif18a, and HSET; and chromosomes defined by DAPI signal on sum projections for Kid.

#### Quantification of polar kinetochore pairs

Polar kinetochores in all experiments were defined as kinetochore pairs that are closer to one of the spindle poles than to the equatorial plane. In confocal-based experiments polar chromosomes were defined from the onset of imaging. The number of polar kinetochore pairs in cells that were imaged from the onset of mitosis, mainly related to LLSM-based experiments, was quantified 12 minutes from the onset of mitosis and then every 6 minutes until the onset of anaphase or until the end of the imaging on maximum intensity projections of all acquired z planes.

To calculate the distance between kinetochores in fixed samples, two points were placed at the center of the signal in each pair of kinetochores using a ‘Point tool’ in ImageJ. Congression velocity was calculated for each pair of kinetochores in the last 6 minutes before the center of the pair of kinetochores surpassed 2 µm from the equatorial plane measured as the nearest distance from a center of a pair to a plane. These times represent the fast kinetochore movement towards the plate. Aligned kinetochore pair was defined as every pair that was found 3 μm from the equatorial plane at any given time. The equatorial or metaphase plane was defined in each time frame as the line perpendicular to the line connecting the centrosomes at their midpoint. The angle between the kinetochore pairs and the main spindle axis was defined as an angle between a line connecting two centrosomes and a line connecting the center of the kinetochore pair and a spindle pole nearest to the respective pair.

In images acquired using STED microscopy, lateral attachments were defined as those in which the kinetochore did not form visible attachments with microtubules that terminate at the kinetochore, defined by the CENP-A signal; end-on attachments were defined as those in which the kinetochore formed visible attachments with microtubules that terminated at the kinetochore, defined by the CENP-A signal, with no visible microtubule signal just below or above the kinetochore. When scoring successful alignment of polar chromosomes, alignment was considered successful if a chromosome, initially polar at the beginning of imaging, moved to within 2 μm of the equatorial plane after 30 minutes. The 30-minute period was chosen because it represents the minimum imaging time used for the confocal-based microscopy assay, and it covers the majority of the imaged cells.

The alignment categories were defined as follows: “No metaphase plate” indicated that no metaphase plate is formed. “High misalignment” indicated a metaphase plate is formed, but more than 5 chromosomes are outside the plate. “Low misalignment” indicated a situation where a metaphase plate is formed, with fewer than 5 chromosomes outside the plate. Finally, “Tight metaphase plate” signified the formation of a tight metaphase plate with no chromosome misalignment. “Maximum spread of chromosomes” was measured on maximum intensity projections as the nearest distance between centers of two most separated kinetochore pairs.

#### Image processing and statistical analysis

Image processing was performed in ImageJ (National Institutes of Health, Bethesda, MD, USA). Quantification and statistical analysis were performed in MatLab. The figures were assembled in Adobe Illustrator CS6 (Adobe Systems, Mountain View, CA, USA). Raw images were used for quantification. The images of spindles were rotated in every frame to fit the long axis of the spindle to be parallel with the central long axis of the box in ImageJ and the spindle short axis to be parallel with the central short axis of the designated box in ImageJ. The designated box sizes were cut to the same dimensions for all panels in the figures where the same experimental setups were used across the treatments. When comparing different treatments in channels in which the same protein was labeled, the minimum and maximum intensity of that channel was set to the values in the control treatment. When indicated, the smoothing of the images was performed using the “Gaussian blur” function in ImageJ (s=0.5-1.0). Colour-coded maximum intensity projections of the z-stacks were done using the “*Temporal colour code*” tool in Fiji by applying “16-color” or “Spectrum” lookup-table (LUT) or other LUT as indicated. For the generation of univariate scatter plots, the open Matlab extension “UnivarScatter” was used.

Data are given as mean ± standard deviation (s.t.d.), unless otherwise stated. The mean line was plotted to encompass a minimum of 60% of the data points for each treatment. Other dispersion measures used are defined in their respective figure captions or in Fig. 1 and Extended Data Fig. 1 if the same measures are used across all figures. The exact values of n are given in the respective figure captions, where n represents the number of cells or the number of tracked kinetochore pairs, as defined for each n in the figure captions or tables. The number of independent biological replicates is also given in the figure captions. An independent experiment for acute drug treatments was defined by the separate addition of a drug to a population of cells in a dish. The number of cells imaged simultaneously ranged from 1 to 7, depending on the specific microscopy system used. The p values when comparing data from multiple classes that followed a normal distribution were obtained using the one-way ANOVA test followed by Two-sided Tukey’s Honest Significant Difference (HSD) test (significance level was 5%). p < 0.05 was considered statistically significant, very significant if 0.001 < p < 0.01 and extremely significant if p < 0.001. Values of all significant differences are given with the degree of significance indicated (∗0.01 <p < 0.05, ∗∗0.001 < p < 0.01, ∗∗∗p < 0.001, ∗∗∗∗ < 0.0001). For the linear regression correlation measure between two parameters, the nonparametric Spearman correlation coefficient, termed rs, was used where p<0.001, calculated using the ‘corr’ function in Matlab (Statistics Toolbox R14). No statistical methods were used to predetermine the sample size. The experiments were not randomized and, except where stated, the investigators were not blinded to allocation during experiments and outcome evaluation.

## DATA AVAILABILITY

Source data for the main figures is provided with this paper. All other data supporting the findings of this study are available from the corresponding authors on reasonable request.

## CODE AVAILABILITY

Codes used to analyze and plot the data are available from the corresponding authors on request.

## EXTENDED DATA FIGURE LEGENDS

**Extended Data Fig. 1.**
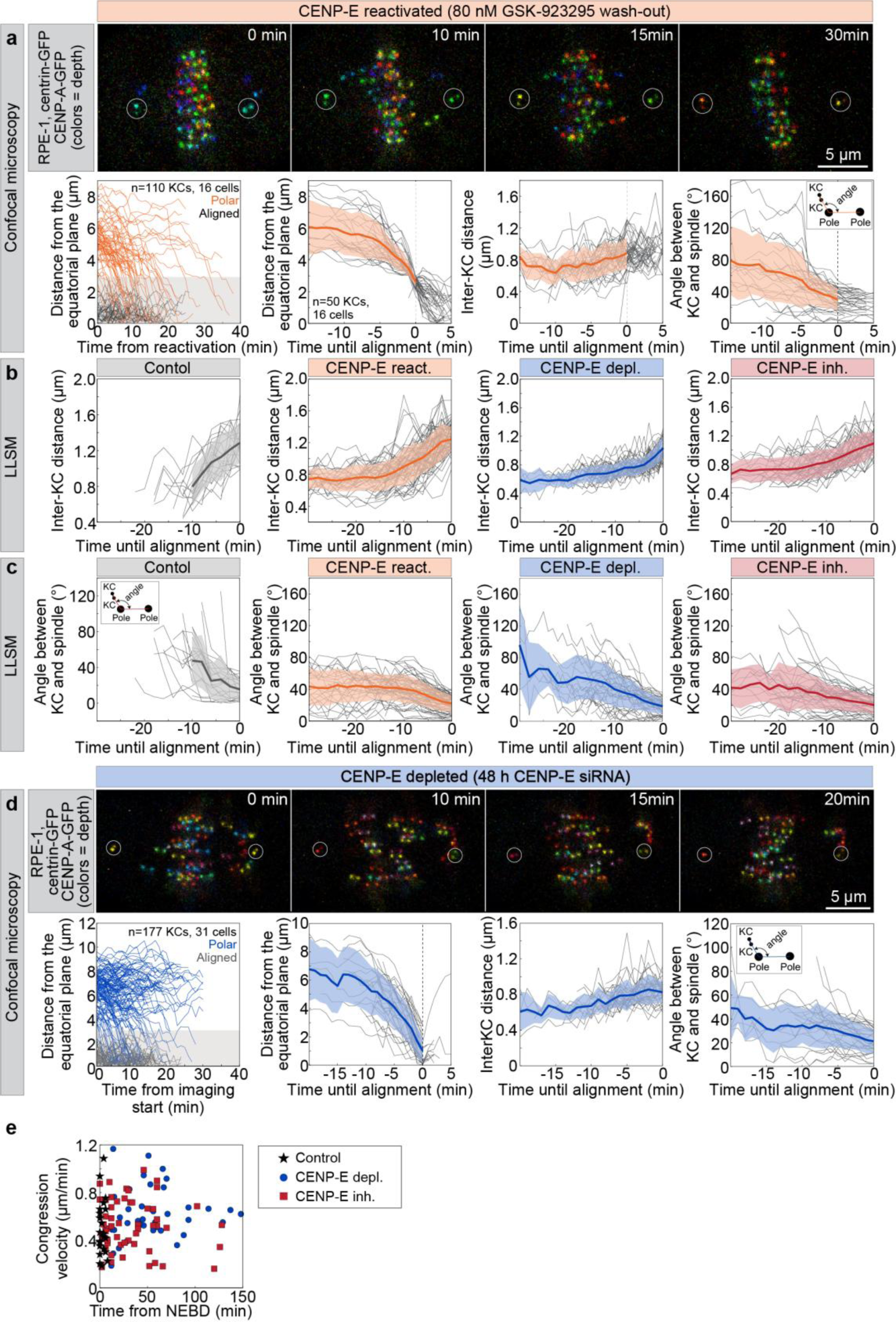
Chromosome movement during congression in RPE-1 cells is similar regardless of CENP-E activity or mitotic duration. **(a)** Representative confocal images of an RPE-1 cell expressing CENP-A-GFP and centrin1-GFP after washout of 80 nM CENP-E inhibitor GSK923295, shown as maximum intensity projections color-coded by depth (color bar in Fig. 1a). Plots show the distance from the equatorial plane (top), interkinetochore distance (inter-KC, middle), and angle between sister kinetochores and the spindle axis, as illustrated in the top-left scheme (bottom), of sister kinetochores for initially polar (orange) pairs, over time from CENP-E inhibitor washout, with means (thick lines) and standard deviations (grey shading). Centrioles are marked by white circles. In top left graph black lines represent aligned kinetochores at the moment of inhibitor washout. **(b)** Distance between polar kinetochores (inter-KC) until alignment across indicated treatments imaged by lattice light sheet microscopy (LLSM). **(c)** Angle between polar kinetochores and the spindle axis, as depicted in the top-left scheme, until alignment across treatments imaged by LLSM, (*d*) Representative confocal images of an RPE-1 cell expressing CENP-A-GFP and centrin1-GFP after 48h CENP-E depletion, with similar plots for kinetochore distances, inter-KC, and angles as in (a). (**e**) Congression velocity of polar kinetochore pairs during the last 6 minutes of chromosome congression plotted against time from nuclear envelope breakdown (NEBD) when congression event was initiated, and for treatments indicated. Abbreviations: KC, kinetochore; depl., depleted; inh., inhibited; react., reactivated; NEBD, nuclear envelope breakdown.

**Extended data Fig. 2.**
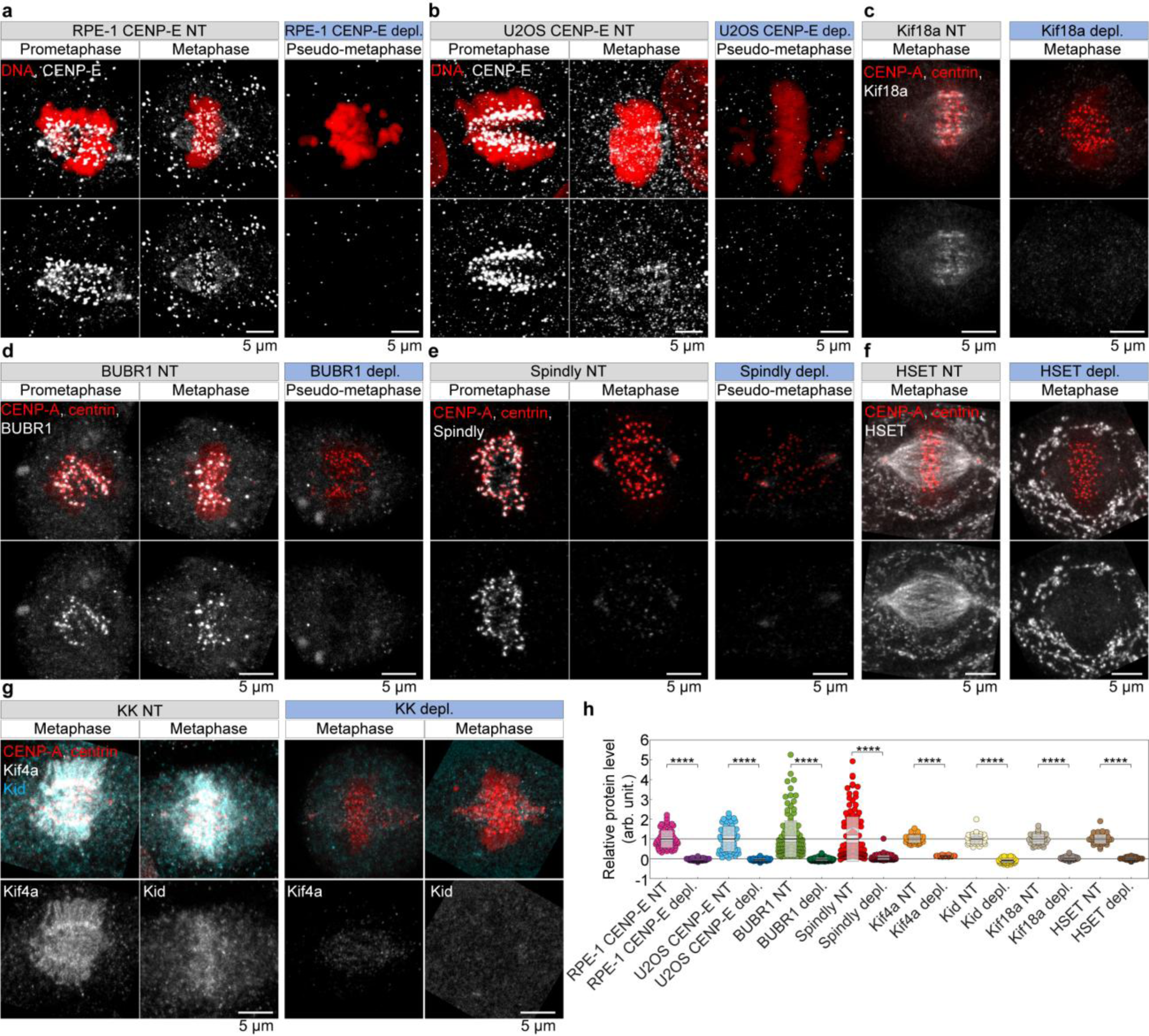
Immunofluorescence-based assessment of RNA interference efficiency across multiple targets. **(a–g)** Representative images of RPE-1 (a, c–g) and U2OS (b) cells treated with non-targeting (NT, grey boxes) or targeting siRNAs (blue boxes), immunostained with antibodies against the indicated proteins. Images show different mitotic stages as labeled. Panels on the left correspond to NT siRNA-treated cells, while panels on the right show cells treated with siRNAs targeting the specific proteins indicated. RPE-1 cells were either wild-type and stained with DAPI (a, red) or expressed CENP-A–GFP and centrin–GFP (c–g, red). U2OS cells were wild-type and stained with DAPI (b, red). Top panels show maximum intensity projections of merged fluorescence channels, and bottom panels show grayscale images of antibody signal for the targeted protein only. **(h)** Quantification of average protein levels normalized to the NT control group. Sample sizes were: 70, 66, 56, 53 cells for CENP-E; 137, 116, 121, 144 kinetochores from 14, 13, 15, and 15 cells for BubR1 and Spindly; and 36, 42, 35, 43, 34, 38, 42, and 36 cells for Kid through HSET, all from at least two independent biological replicates. Statistical analysis was performed using ANOVA with post-hoc Tukey’s HSD test. Symbols indicate: ****p < 0.0001; NT, non-targeting; depl., depleted; KK, Kid and Kif4a.

**Extended Data Fig. 3.**
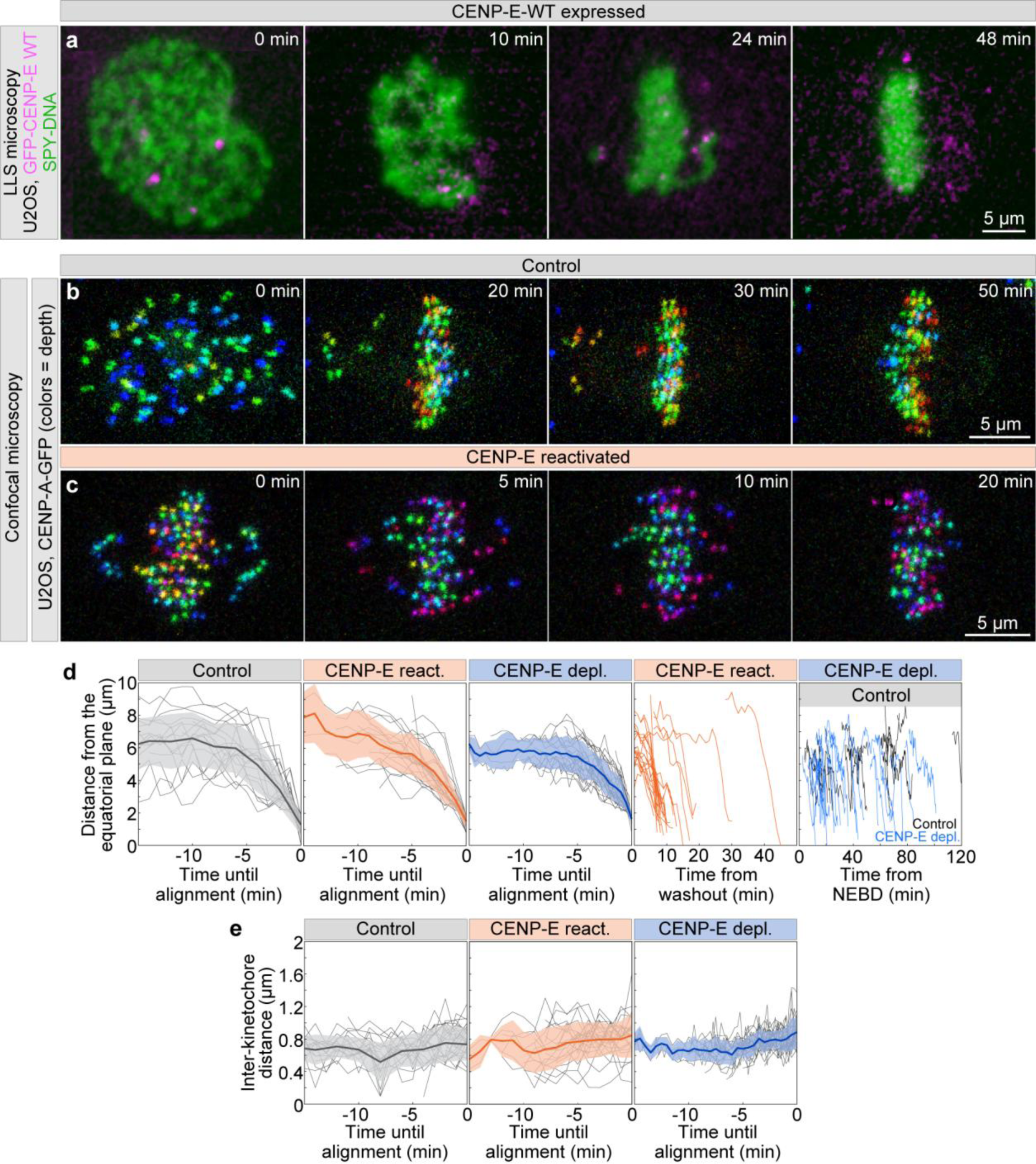
Chromosome movement during congression is similar in U2OS cells despite varying CENP-E activity. **(a)** Representative image of a U2OS cell expressing GFP– CENP-E wild-type (WT) (magenta) following depletion of endogenous CENP-E, stained with 10 nM SPY650–DNA (green). Time 0 corresponds to the onset of nuclear envelope breakdown (NEBD). **(b, c)** Representative examples of U2OS cells expressing CENP-A–GFP imaged by confocal microscopy at different time points in a control DMSO-treated cell (top) and after washout of 200 nM GSK923295. Time 0 corresponds to the onset of NEBD in (a) or the time of inhibitor washout in (b). Images are shown as maximum intensity projections and color-coded by depth, as indicated by the color bar in Fig. 1a. **(d, e)** Quantification of the distance from polar kinetochores to the spindle equatorial plate (d) and interkinetochore distance (e) over indicated time points, representing time until alignment, time from inhibitor washout, or time from NEBD, as shown, under different treatment conditions indicated above each graph. Sample sizes for cells and kinetochores are provided in Fig. 1. Abbreviations: react., reactivated; depl., depleted.

**Extended Data Fig. 4.**
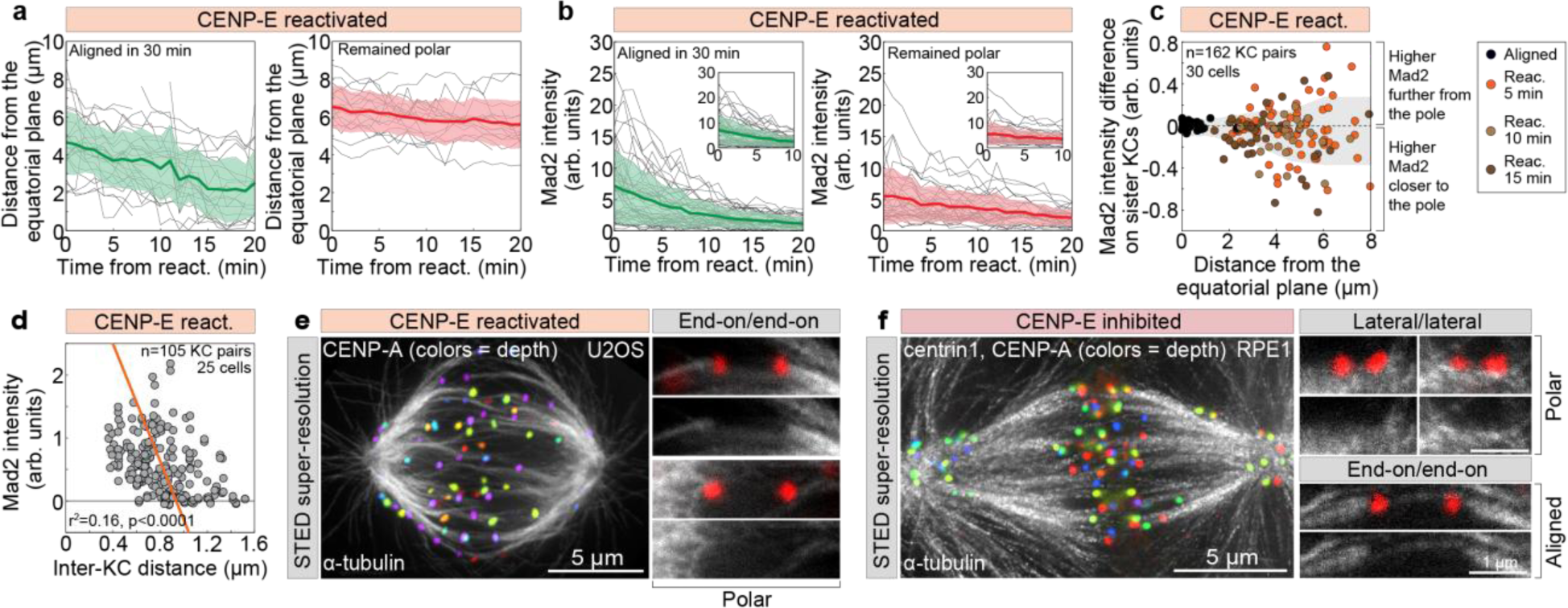
The Mad2 signal on polar kinetochores gradually decreases before the onset of congression movement. **(a, b)** Distance from the equatorial plane (a) and mean Mad2 signal intensity (b) of polar kinetochore pairs over time from washout of the CENP-E inhibitor, shown separately for kinetochore pairs that aligned within 30 minutes (green, left) and pairs that did not align in the same period (red, right). **(c)** Distance of kinetochore pairs from the equatorial plane plotted against the difference in Mad2 levels between sister kinetochores based on their proximity to the nearest pole under the indicated conditions (right). **(d)** Interkinetochore (inter-KC) distance of sister kinetochore pairs versus mean Mad2 signal intensity with linear regression (line). **(e, f)** Representative spindles from a U2OS cell 8 minutes after CENP-E reactivation (e) and an RPE-1 cell after continuous CENP-E inhibition (f), immunostained for α-tubulin (grey) and imaged by stimulated emission depletion (STED) microscopy. U2OS cells expressed CENP-A-GFP and RPE-1 cells expressed CENP-A-GFP and centrin1-GFP, which are color coded by depth, as specified by the color bar in Fig. 1a (left). Images are maximum intensity projections. Insets show kinetochore pairs with microtubules from different attachment categories as depicted, alongside schematic representations of microtubules (white) and kinetochores (red) for the respective insets (right). Abbreviations: react., reactivation; KC, kinetochore; inh., inhibited.

**Extended Data Fig. 5.**
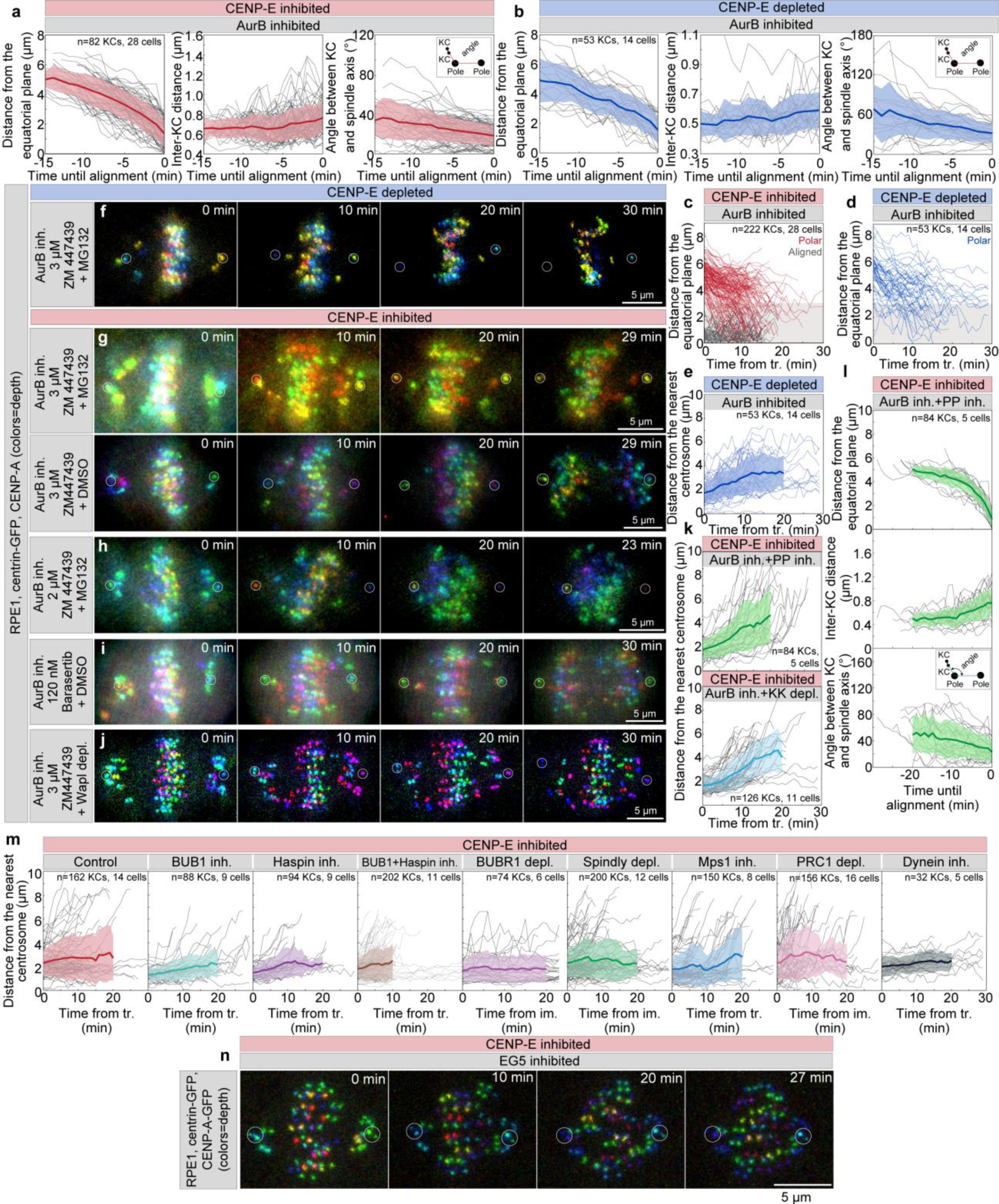
Chromosome congression without CENP-E is mainly limited by kinetochore-localized Aurora. **B**. **(a, b)** Distance to the equatorial plane (left), interkinetochore distance (middle), and angle between sister kinetochores and spindle axis (right, scheme) over time until alignment after acute Aurora B inhibition (3 µM ZM447439) in CENP-E inhibited (a) or depleted (b) cells. Thick lines: means; grey areas: SD. Centrioles circled in white. **(c–e)** Distance to equatorial plane of initially polar (red/blue) and aligned (black) sister kinetochores over time from Aurora B inhibitor addition in CENP-E inhibited (c) and depleted (d) cells; (e) distance to nearest spindle pole of polar kinetochores in CENP-E depleted cells. **(f–j)** Representative RPE1 cells expressing CENP-A-GFP and centrin1-GFP after treatments: acute Aurora B inhibition with 3 µM ZM447439 in CENP-E depleted (f), in CENP-E inhibited cells ± 20 µM MG132 (g), Aurora B inhibition with 2 µM ZM447439 + MG132 (h), Aurora B inhibition by 120 nM barasertib + DMSO (i), and Aurora B inhibition with ZM447439 + DMSO in WAPL-depleted, CENP-E inhibited cells (j). Centrioles circled. **(k)** Distance to nearest centrosome of polar kinetochores over time for indicated treatments. **(l)** Distance to equatorial plane (top), inter-KC distance (middle), and angle to spindle axis (bottom) in cells with combined Aurora B (3 µM ZM447439) and protein phosphatase (1 µM Okadaic acid) inhibition under CENP-E inhibition. (**m**) Distance to the nearest centrosome over time for initially polar chromosomes in different treatments. **(n)** Example of RPE1 cell after acute addition of 100 µM Eg5 inhibitor monastrol in CENP-E inhibited cells. Time indicated from treatment. All images are maximum projections color-coded by depth (Fig. 1a color bar). Abbreviations: KC, kinetochore; depl., depleted; inh., inhibited; react., reactivated; tr., treatment; im., imaging; KK, Kid and Kif4a; PP, protein phosphatases.

**Extended Data Fig. 6.**
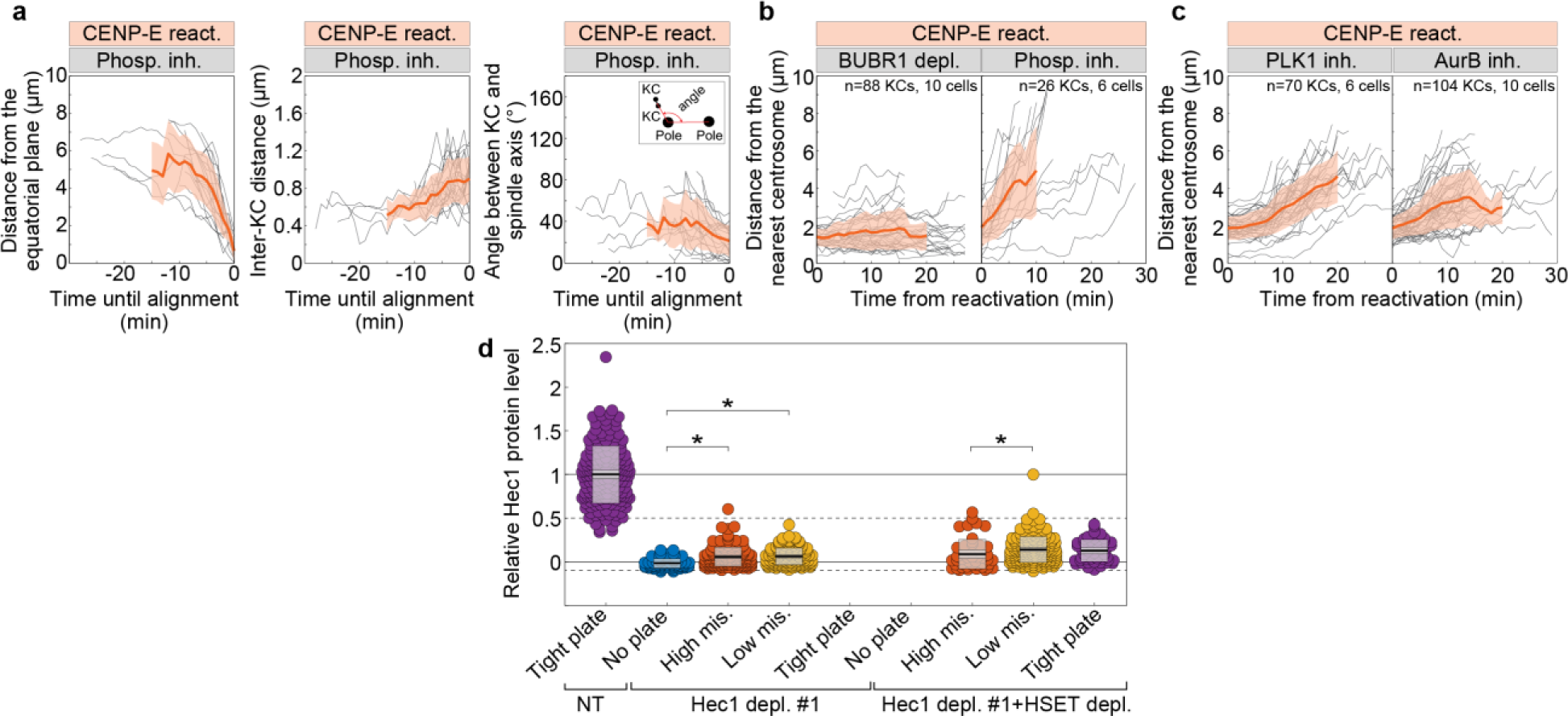
Chromosome congression after CENP-E reactivation is independent of protein phosphatase activity. **(a)** Distance to the equatorial plane of initially polar kinetochore pairs (left), interkinetochore distance (middle), and angle between sister kinetochores and the spindle axis (right, scheme) over time until alignment in cells treated with 1 µM Okadaic acid to acutely inhibit protein phosphatases after CENP-E inhibitor washout. Thick lines: means; grey areas: SD. **(b, c)** Distance to nearest centrosome of initially polar kinetochore pairs over time from treatments indicated above each graph. **(d)** Average Hec1 levels on kinetochores normalized to non-targeting (NT) controls across treatments and alignment categories as indicated. Statistics: ANOVA with post-hoc Tukey’s HSD test. Symbols: * P ≤ 0.05. Abbreviations: KC, kinetochore; inh., inhibited; depl., depleted; react., reactivated; mis., misalignment.

## VIDEO LEGENDS

**Video 1. Lattice-light sheet microscopy (LLSM)-based assay for large-scale live cell imaging.** Dozens of RPE-1 cells stably expressing CENP-A-GFP and Centrin1-GFP (16-colors LUT) treated with DMSO (control) were imaged by lattice light sheet microscopy (LLSM) immediately after drug addition. Kinetochores and centrioles are color-coded by depth from blue to red.

**Video 2. RPE1 cells stably expressing CENP-A-GFP and Centrin1-GFP in control and CENP-E reactivated conditions, imaged by LLSM.** RPE-1 cells stably expressing CENP-A-GFP and Centrin1-GFP (16-colors LUT) treated with DMSO (control, left) or after washout of the CENP-E inhibitor (CENP-E reactivated, right) were imaged immediately following drug addition (left) or drug washout (right). Kinetochores and centrioles are color-coded by depth from blue to red using the 16 Colors LUT.

**Video 3. RPE1 cells stably expressing CENP-A-GFP and Centrin1-GFP in CENP-E depleted and CENP-E inhibited conditions, imaged by LLSM.** RPE-1 cells stably expressing CENP-A-GFP and Centrin1-GFP (16-colors LUT) treated with CENP-E siRNA for 48 h (CENP-E depleted, left) or with CENP-E inhibitor for 3 h (CENP-E inhibited, right) were imaged at the respective time points. Kinetochores and centrioles are color-coded by depth from blue to red using the 16 Colors LUT.

**Video 4. RPE1 cells stably expressing CENP-A-mCerulean and Mad2-mRuby following reactivation of CENP-E activity, imaged by confocal microscopy.** RPE-1 cell stably expressing CENP-A-mCerulean (grey) and Mad2-mRuby (red), treated with CENP-E inhibitor for 3 hours followed by drug wash-out, was imaged immediately after wash-out. Scale bar, 5 μm.

**Video 5. RPE1 cells stably expressing CENP-A-GFP and Centrin1-GFP in CENP-E inhibited condition following addition of inhibitors of Aurora kinases, and in CENP-E reactivated condition following depletion of BubR1 and addition of inhibitor of phosphatases, imaged by confocal microscopy.** RPE-1 cells stably expressing CENP-A-GFP and Centrin1-GFP (16-colors LUT) treated with CENP-E inhibitor for 3 hours followed by acute inhibition of Aurora B kinase (top left) or Aurora A kinase (top right), and cells treated with CENP-E inhibitor for 3 hours followed by drug wash-out in BubR1-depleted cells (bottom left) or acute phosphatase inhibition (bottom right). Cells were imaged immediately after the last drug addition or wash-out. Kinetochores and centrioles are color-coded for depth from blue to red using the 16 Colors LUT, with noise reduction applied via the Despeckle function in ImageJ.

